# Coordinated IFN-γ/TNF Axis Drives Selective Loss of Activated Enteric Glia in Inflammatory Bowel Diseases

**DOI:** 10.1101/2025.04.03.646820

**Authors:** Marvin Bubeck, Klara A. Penkert, Heidi Limberger, Miguel González Acera, Christina Plattner, Svenja Ziegler, Anoohya Muppirala, Patrycja Forster, Manuel Jakob, Reyes Gamez-Belmonte, Lena Erkert, Subhash Kulkarni, Claudia Günther, Raja Atreya, Anja A. Kühl, Ahmed N. Hegazy, Kai Hildner, Zlatko Trajanoski, Britta Siegmund, Markus F. Neurath, IBDome Consortium, Meenakshi Rao, Fränze Progatzky, Dieter Chichung Lie, Christoph Becker, Chiara Romagnani, Leif S. Ludwig, Christoph S. N. Klose, Jay V. Patankar

**Affiliations:** Department of Medicine1, Universitätsklinikum Erlangen and Friedrich-Alexander-Universität Erlangen-Nürnberg (FAU), Erlangen, Germany; Deutsches Zentrum Immuntherapie (DZI), Erlangen, Germany; Berlin Institute of Health at Charité – Universitätsmedizin Berlin, Berlin, Germany; Max-Delbrück-Center for Molecular Medicine in the Helmholtz Association (MDC), Berlin, Institute for Medical Systems Biology (BIMSB), Berlin, Germany; Charité – Universitätsmedizin Berlin, Berlin, Germany; Biocenter, Institute of Bioinformatics, Medical University of Innsbruck, Innsbruck, Austria; Charité, Department of Hematology, Oncology, and Tumor Immunology; Division of Gastroenterology, Department of Pediatrics, Boston Children’s Hospital and Harvard Medical School, Boston, United States; Charité – Universitätsmedizin Berlin, corporate member of Freie Universität Berlin and Humboldt-Universität zu Berlin, Department of Microbiology, Infectious Diseases and Immunology, Berlin, Germany; Division of Gastroenterology, Dept of Medicine, Beth Israel Deaconess Medical Center, Boston, MA; Division of Medical Sciences, Harvard Medical School, Boston, MA; Charité – Universitätsmedizin Berlin, Freie Universität Berlin, corporate member of Humboldt Universität zu Berlin, iPATH.Berlin, Berlin, Germany; Charité-Universitätsmedizin Berlin, corporate member of Freie Universität Berlin and Humboldt-Universität zu Berlin, Department of Gastroenterology, Infectious Diseases and Rheumatology, Berlin, Germany; The IBDome Consortium; The Kennedy Institute of Rheumatology, University of Oxford, Oxford, United Kingdom; Institut für Anatomie, Lehrstuhl für Mikroskopische Anatomie und Molekulare Bildgebung, Friedrich-Alexander-Universität Erlangen-Nürnberg (FAU), Erlangen, Germany; Institute of Medical Immunology, Charité Universitätsmedizin Berlin, Corporate Member of Freie Universität Berlin and Humboldt Universität zu Berlin, Berlin, Germany; Innate Immunity, Deutsches Rheuma-Forschungszentrum Berlin (DRFZ), ein Leibniz Institut, Berlin, Germany

**Author notes:** Corresponding author Correspondence: Jay V. Patankar, PhD Universitätsklinikum Erlangen, Department of Medicine1, Gastroenterology, Pneumology, Endocrinology Erlangen, Germany.

**Keywords:** Inflammatory bowel disease, activated enteric glial cells, single nucleus mRNA Sequencing, necroptosis, gut motility

## Abstract

**Background:** Enteric glial cells (EGC) play a crucial role in maintaining gut homeostasis, but their dysregulation in inflammatory bowel diseases (IBD) remains poorly understood. Emerging preclinical data suggests activated EGC have beneficial roles in controlling gut pathophysiology.

**Objective:** Understanding EGC activation and adaptation during experimental and clinical IBD.

**Design:** We provide the first highly integrated approach to identify EGC activation signature in IBD. Profiling 390 samples from IBD patients via bulk and single-nucleus (sn) transcriptomics and replicate the findings on publicly available bulk and single-cell (sc) datasets from 1160 patients and 19,000 single EGC. Preclinical modelling of Th1/Th17 inflammation, reporter-assisted EGC sorting, analysis of regulated cell death, and *Casp8* ablation in EGC was performed

**Results:** We identified novel IBD type and sampling associated EGC activation signature. Specific EGC activation markers were shared in biopsies and resection specimens, and were divergent between Crohn’s disease and Ulcerative colitis. Preclinical modelling of intestinal inflammation identified combinatorial TNF and IFN-γ-driven activation of EGC, associated with elevated necroptosis, and negatively impacting gut motility. Genetic-reporter-enabled sorting and downstream analyses confirmed TNF and IFN-γ-driven EGC necroptosis, potentiated by *Casp8* deficiency. Furthermore, snRNA-Seq from IBD patient samples confirmed elevated cell death signature in activated but not in rare neuroglia progenitor-like cluster.

**Conclusion:** Our findings identify IBD type-associated activated EGC markers involved in immune and epithelial homeoastasis. We uncover necroptosis of activated EGCs as a constituent of intestinal inflammation. Advancing our understanding of activated EGC survival is pivotal in elucidating their complex roles in maintaining gut immune-epithelial homeostasis.

**What is already known on this topic:** Activated EGC have emerged as important contributors in maintaining epithelial, immune and neuronal homeostasis. Increasing evidence from mouse studies points to the role of activated EGC in epithelial regeneration, tolerogenic T-cell activation, relaying psychological stress to the enteric nervous system, post-injury neurogenesis, and helminth clearance. Nevertheless, no consensus has emerged on what might define activated EGC in the context of IBD and how EGC turnover is affected in gut inflammation, limiting translation of their disease associated roles.

**What this study adds:** By combining bulk with single cell and single nucleus transcriptomes from IBD patients we identified new IBD type– and location-associated EGC activation signatures. Some of these are conserved with mouse EGC in gut inflammation models. We identified osteopontin an immunomodulator and *Wnt6* an epithelial morphogen elevated in IBD EGC. We also identified IBD-associated EGC cell clusters, which display higher expression of cell death pathway transcripts. To investigate EGC turnover, we utilized preclinical models and found rapid EGC activation upon Th1/Th17 inflammation. This was associated with elevated EGC activation and caspase-independent necroptotic cell death. *Ex vivo* experiments showed a combinatorial requirement of IFN-γ and TNF in mediating EGC necroptosis. Our findings were replicated on multiple publicly available sc-RNA sequencing datasets from IBD patients.

**How this study might affect research, practice or policy:** Expanding on the available repertoire of EGC activation markers in IBD, both shared and unique to sampling procedure, disease type, and location will provide researchers with tools to identify EGC homeostasis during IBD. Moreover, the nature of the identified markers will stimulate research into specific EGC pathways triggered in inflammation. Adding to this, the rapid induction in pathological death of activated but not naive EGC upon IFN-γ and TNF stimulation will shed light on EGC adaptation and turnover. Our identification of markers of activated EGC with immuno-modulatory and epithelial-regenerative properties, including osteopontin and wingless family of morphogenes will stimulate further research in EGC-immune and EGC-epithelial communication in the context of IBD.

## Introduction

Inflammatory bowel diseases (IBD) pose a chronic inflammatory challenge, demanding concerted actions of multiple cells in restoring homeostasis. Enteric glial cells (EGC) display distinct morphotypes and activation states controlling gut motility, visceral hypersensitivity, and lymphocyte activation [1, 2, 3, 4, 5]. Data from preclinical models has revealed that activated EGC subtypes control epithelial, immune, and neuronal homeostasis during intestinal inflammation. For example, during intestinal inflammation activated EGC express: 1. MHC-II, inducing tolerogenic T-cells [6], 2. Wnt ligands regulating epithelial regeneration [3], 3. CSF1 promoting monocyte recruitment [2], and 4. CXCL10 promoting helminth clearance [7].

Despite this, the translation of these findings in the context of IBD has been limited largely due to a lack of consensus on what constitutes activated EGC. Through multiple studies in mouse models GFAP has emerged as a bona fide activation marker for EGC. However, its expression and specificity in human EGC is highly contested. Furthermore, the status of S100B, another prevailing activation marker of human EGC, was also challenged recently [8]. The S100B regulates survival and can act as a damage-associated molecular pattern and hence its elevation has been linked with EGC activation. Another broadly expressed maker for EGC is PLP1, with established role in myelination in the peripheral nervous system. However, its regulation during gut inflammation and its functions in the non-myelinated ENS are poorly defined with recent studies pointing to a role in the regulation of gut motility in mice [4]. This is reflected in the disparity in previous reports, some indicating EGC elevation, whereas other showing EGC apoptosis and reduction in EGC frequency in IBD [9, 10, 11, 12, 13, 14]. This is fuelled by challenges in analysing all EGC subsets from patient samples, since translational research is dominated by data derived from mucosal biopsies, which miss out on myenteric and longitudinal muscle EGC, limiting a clearer insight on EGC activation markers in IBD.

An early consequence of gut inflammation is dysmotility, induced in acute conditions and sustained by dysbiotic microbial insults. This can trigger enteric neuronal pyroptosis and, as we have recently shown, neuronal, but not EGC ferroptosis [15, 16, 17]. It is currently unknown whether and how EGC may evade programs of cytokine-mediated pathological cell death such as necroptosis and how this affects EGC turnover in IBD. This is especially important in the context of anti-TNF therapy, TNF being an upstream inducer of necroptosis, where few studies in experimental colitis have investigated neuroglia protection after anti-TNF treatment [18].

Based on these considerations, we took a highly integrated approach to first identify EGC activation markers in IBD types, comparing biopsy and resection tissues in bulk, single nucleus (sn), and single cell (sc) transcriptomes from IBD patients. We uncovered that specifically the activated EGC subsets display elevated expression of cell death pathway signatures. Analysis of sorted EGC revealed a combined requirement of TNF and IFN-γ in mediating EGC necroptosis. Our study provides novel EGC-specific activation markers elevated in IBD, which regulate EGC-immune and EGC-epithelial homeostasis and highlight TNF and IFN-γ mediated activated EGC necroptosis as a component feature in Th1/Th17-driven intestinal inflammation. Further studies aimed at investigating EGC turnover, cellular states, and EGC –immune and –epithelial crosstalk are necessary for teasing out the translational impact of crucial EGC functions in IBD.

## Results

### Identification of EGC activation markers across IBD

An initial analysis of resection and mucosal biopsy data from IBD patient transcriptomes (IBDome cohort) and previous microarray datasets from IBD patients revealed a divergent expression of the canonical EGC markers *S100B* and *PLP1* (Supplemental Fig 1A-D). To identify IBD-associated EGC-specific markers, we leveraged the scIBD atlas of over 1.1 million IBD sc-transcriptomes. Comparing these sc expression profiles of EGC versus all other cells, top IBD-associated EGC markers were extracted and tracked across the bulk transcriptomes of our IBDome patient cohort and the Mount Sinai Crohn’s and Colitis Registry (MSCCR). This enabled comparison of IBD-associated, EGC-specific markers across disease types, tissues, and sampling procedures (Fig. 1A) [19, 20].

**Figure 1.**
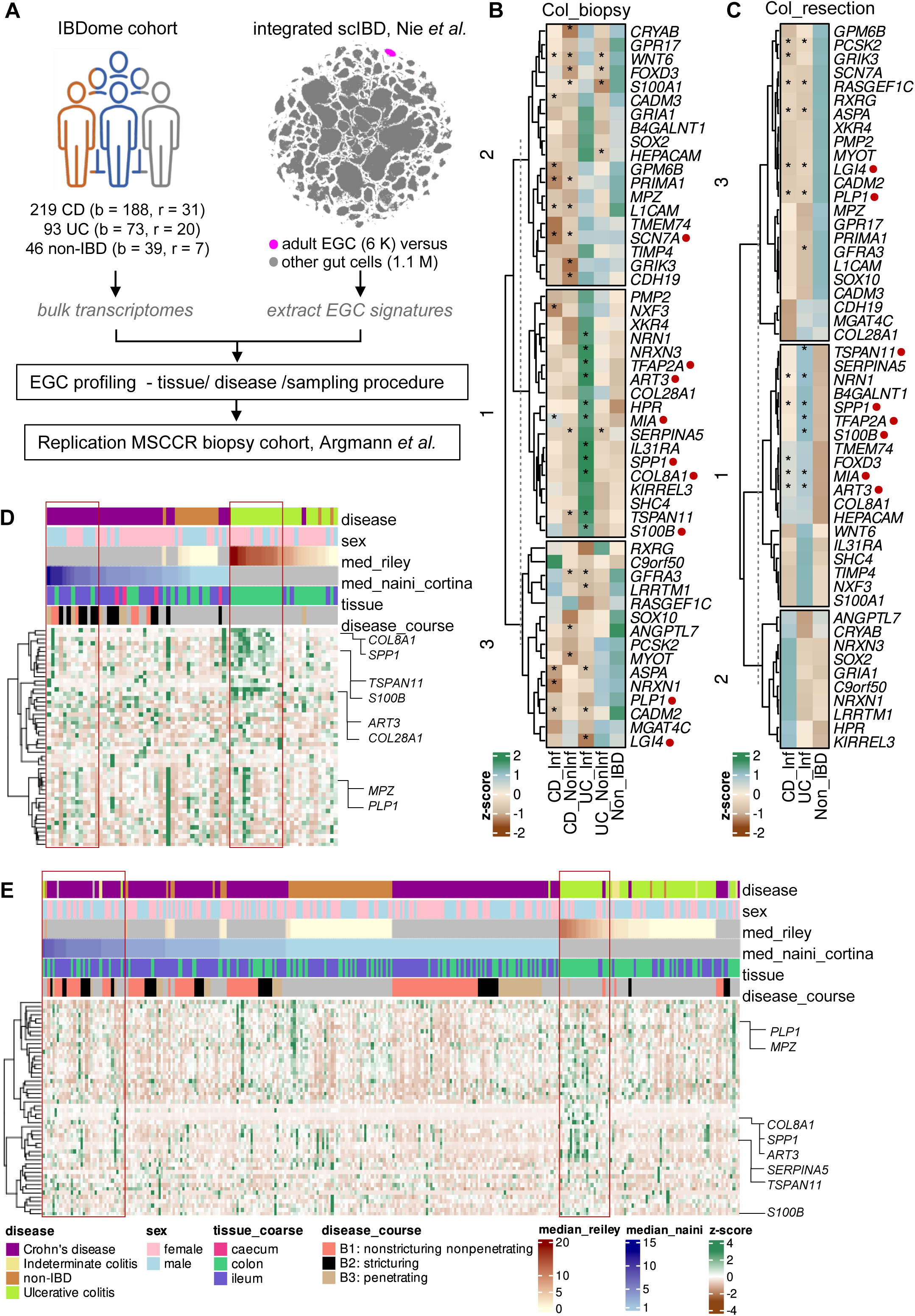
IBD subtype, sampling procedure, and inflammation-associated enteric glia signatures. **A)** Scheme depicting the composition of the IBDome cohort with number of patients split by sampling procedure (b = biopsy, r = resection), the extraction of enteric glial cell (EGC) enriched gene signature from the integrated single cell IBD transcriptomes [20] and comparison and replication strategy on the Mount Sinai Crohn’s and Colitis Registry cohort [19]. **B & C)** Heatmaps comparing mean z-scores calculated from normalized counts for indicated EGC genes across IBD samples from the IBDome cohort from colonic (Col_) biopsies (B) or resections (C) with red dots indicating genes of interest. Asterisk indicate p-values < 0.05 from Mann-Whitney test applied on normalized counts against the Non_IBD group. **D & E)** Heatmaps showing sample-wise z-scores for the EGC enriched genes as in (B) & (C) with top annotations sorted by descending order for the median (med_) histologic severity scores by IBD type from resections (D) and biopsies (E).

Based on their expression across all samples included in our IBDome cohort, correlations could be established not only between the expression of the identified EGC markers in a given sample, but also with the respective histological inflammation scores for that sample. For example, this could be visualized for colonic biopsy samples as a correlation network identifying new EGC activation markers *SPP1, MIA,* and *ART3* that correlated well with the Riley score for UC and to some degree with the Naini Cortana score for CD, but poorly with naïve/ canonical EGC-specific markers such as *PLP1* (Supplemental Fig. 1E).

Analysis of scaled expression compared against non-IBD control samples, revealed several new inflammation-associated markers in colonic biopsies and resection samples of inflamed UC patients (Fig. 1B, C). The analysis also revealed that a number of these EGC markers associated with inflammation status, irrespective of sampling procedure, implying a conserved regulation in mucosal and myenteric EGC (Fig. 1D = resection, Fig. 1E = biopsy). Among these, *SPP1* and *COL8A1* were upregulated in inflamed UC, but not CD colon biopsies, correlated tightly with modified median Riley scores on the IBDome cohort, were concordant in the resections, and were replicated on the MSCCR cohort Supplemental Fig. 1E, G). Both *ART3* and *MIA* were upregulated in inflamed UC and CD colonic biopsies and resections, and correlated with the canonical activation marker *S100B* (Fig. 1B-C, Supplemental Fig. 1E, G). Whereas, *SCN7A* showed a downregulation that was specific to biopsies from inflamed CD colons in both cohorts (Fig. 1B-C, Supplemental Fig. 1E, G). Interestingly, the increase in *WNT6* was restricted to the colonic inflamed samples and was discordant between the CD and UC samples in the IBDome and MSCCR cohorts, whereas that of *IL31RA* was significant at both tissue locations in biopsies from MSCCR cohort, only reaching significance in the colonic biopsies for the IBDome cohort (Fig. 1B-C, Supplemental Fig. 1F, G).

The overall change in expression seen in the ileal biopsies was more modest than that from the colon biopsies with three genes, namely *S100B*, *SPP1*, and *TFAP2A* being significantly elevated in inflamed samples in both locations from both cohorts (Fig. 1B, Supplemental Fig. 1F-G). The expression of these markers from colon resections showed a significant UC-specific induction (Fig. 1C). The strong UC versus CD disparity in cluster 1 colonic biopsy samples from the IBDome cohort (Fig. 1B) was recapitulated only for *SPP1*, *COL28A1*, *COL8A1*, and *NRXN3* on the MSCCR cohort colonic biopsies (Supplemental Fig. 1G). These findings highlight new inflammation associated EGC markers including the immunomodulatory *SPP1* and *IL31RA* in mucosal and resection samples as well as in CD and UC, and the epithelial morphogen *WNT6* in colonic resection samples.

### Single nuclei RNASeq identifies IBD-associated EGC clusters with elevated cell death signature

So far, we found an induction in specific inflammation-associated EGC activation markers and a slight, but significant reduction in *PLP1* from IBD patient cohorts (Fig. 2B, 1B, Supplemental Fig. 1A-C). This led us to hypothesize that inflammation may trigger a repression in *PLP1* expression or elevated cell death of *PLP1* expressing EGC subsets. To analyse this more directly, we performed sn-RNASeq from full-thickness FFPE tissue sections from the samples in our IBDome patient cohort and non-IBD controls (Supplemental Fig. 2A). After quality control steps, we identified a cluster of 5774 EGC nuclei, identified based on the expression of canonical EGC genes derived from previous literature (Supplemental Fig. 2B) [7, 21, 22, 23, 24]. Sub-clustering of EGC nuclei revealed 9 clusters (Fig. 2A, B). Among these, cluster 4 and 5 were overrepresented in IBD samples with elevated expression of *SRGN* and *CD74* in cluster 4, and *NNMT* and *SOCS3* in cluster 5, resembling reactive and activated glia [25, 26, 27]. Interestingly, the proportion of *PLP1* expressing EGC was slightly lower in these reactive and activated clusters 4 and 5 (Fig. 2B) akin to the slight but significant reduction in *PLP1* expression that we found in the analysis of bulk transcriptomes. This hinted that activated or reactive EGC subsets might be predisposed to elevated cell death and a downregulation of *PLP1*. To analyse whether EGC subtypes may succumb to extrinsic inflammation triggered cell death, we analysed the expression of cell death pathway signature (Supplemental material table ST3), segregated by IBD type and EGC clusters. This revealed a significant elevation in cell death pathway signature in both UC and CD EGCs, specifically showing an elevation in the reactive and activated EGC clusters 4 and 5, which also showed an elevated expression of the chemokine *CCL2* (Fig. 2 C-E).

**Figure 2.**
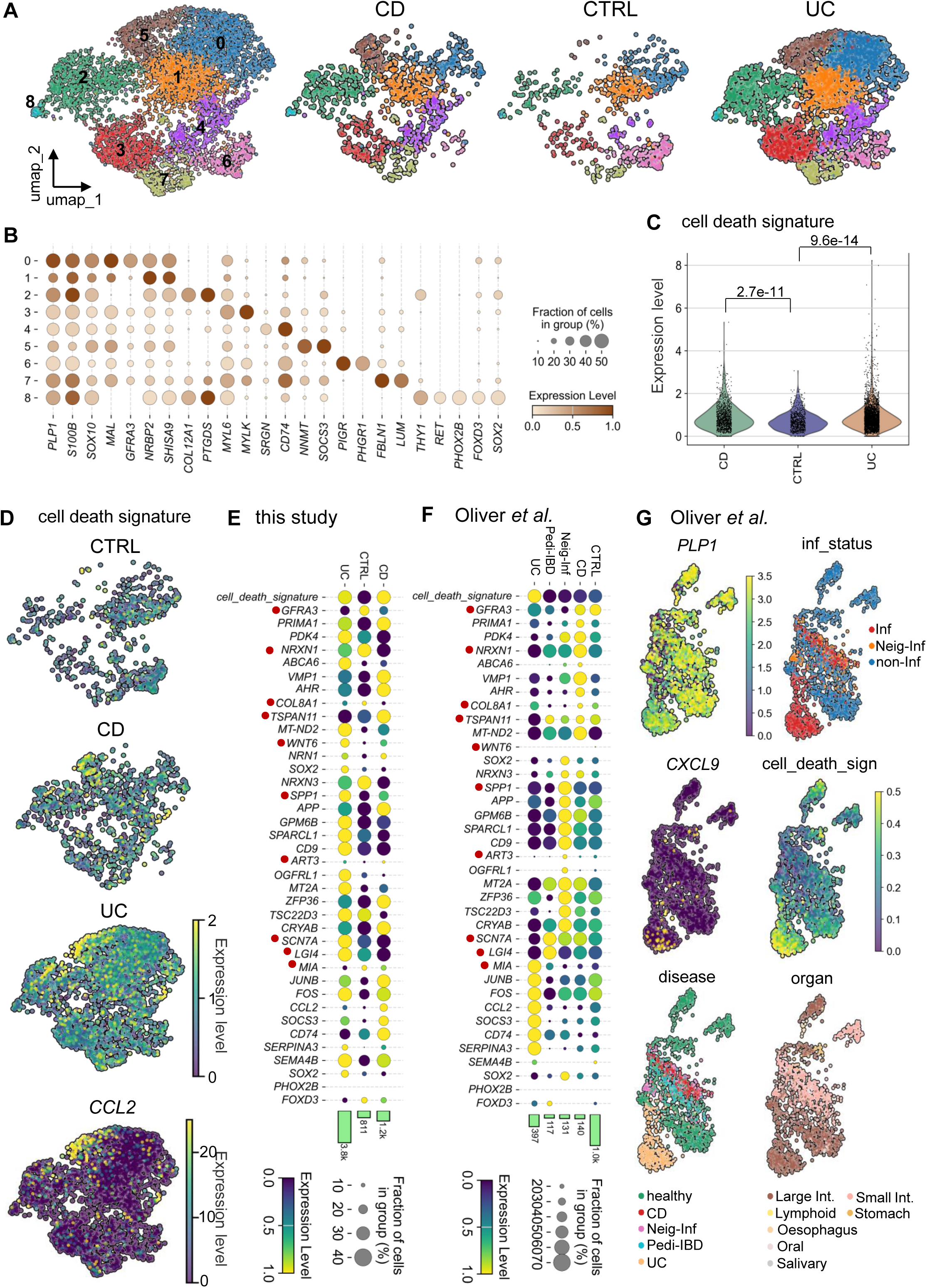
Specific IBD-associated EGC clusters reinforce new activation markers and induction of cell death signature. **A)** Uniform manifold approximation and projection (umap) clustering of mRNA sequencing from single EGC nuclei, split by IBD subtype UC, CD and non-IBD controls (CTRL). **B)** Dot plot showing expression levels of selected canonical EGC and cluster enriched markers. **C)** Violin plots showing quantitative differences in the expression levels of the cell death pathway signature aggregated over all EGC clusters from (A) and segregated by IBD subtype versus CTRL, numbers comparing groups indicate the FDR corrected p-values derived from the two-tailed Mann-Whitney U test. **D)** Scaled average expression of the cell death pathway signature segregated by IBD subtype and CTRL on the umap space. **E&F)** Dot plot showing the expression levels of indicated genes and that of the cell death pathway signature, split by IBD type and CTRL from our snRNA-Seq experiment (E) and from the scRNA-Seq dataset from Oliver *et al*. [28] (F). Bars indicate cell counts per group. The red dots highlight genes of interest identified in Fig. 1. Column abbreviations in F indicate disease type (CD = Crohn’s disease, Neigh-Inf = neighbouring inflamed, Pedi-IBD = paediatric IBD, UC = ulcerative colitis) **G)** umap plots for EGC clusters labelled for gene, or pathway expression, or the cluster annotations for the indicated groups from the single EGC transcriptomes of IBD patients reanalysed from Oliver *et al*. [28]. Annotation abbreviations are as follows inflammation status (inf_status: Inf = inflamed; Neigh-Inf = neighbouring inflamed; non-Inf = non-inflamed), disease type (disease: CD = Crohn’s disease, Neigh-Inf = neighbouring inflamed, Pedi-IBD = paediatric IBD, UC = ulcerative colitis), and organ groups (organ: Int. = intestine).

Apart from these changes, the expression of several IBD-associated EGC-specific markers identified in our analysis earlier (Fig. 1 and Supplemental Fig.1), were recapitulated in our snRNA-Seq data (Fig. 2E). Among these, most prominent were the upregulation of *SPP1* and *WNT6* regulating EGC –immune, –epithelial crosstalk along with the downregulation of *NRXN1* and *GFRA3* regulating EGC –neuron crosstalk (Fig. 2E). In addition, we were also able to detect other genes that were regulated in a disease-specific manner such as *FOS, CCL2*, and *SOCS3* (Fig. 2E). These genes have broad roles in immune homeostasis in many cells and since their expression is not restricted to EGC, they did not appear in the top EGC-specific list in our analysis from Fig 1.

Next, we replicated these findings on 1,797 IBD single EGC transcriptomes reanalysed from Oliver *et al* [28] (Fig. 2E). This dataset has a broader annotation for disease types and we could observe that several of the gene expression patterns detected from our study were concordant with one or the other IBD-related disease types from the Oliver *et al*. dataset (Fig. 2F). This included the elevations in the cell death pathway signature, *SPP1*, *ART3*, *CCL2*, *SOCS3*, and *FOS* among others (Fig. 2F). Interestingly, these changes were accompanied with a slight but significant reduction in *PLP1* expression, specifically in the UC inflamed EGC (Fig. 2G and Supplemental Fig. 2C), which was similar with our bulk transcriptomic analyses (Fig. 1, Supplemental Fig. 1). Furthermore, the same cluster of EGC from inflamed UC samples showed a significant elevation in the expression of the chemokine *CXCL9*, an IFN-γ target gene recapitulating the recent report by Progatzky *et al*. which showed elevated IFN-γ signalling in UC EGC [7]. In addition, this cluster of EGC from inflamed UC samples also showed a significant elevation in the expression of the cell death pathway signature (Fig. 2G and Supplemental Fig. 2C). We further confirmed our findings of this elevated expression of cell death pathway signature, *CXCL9*, and reduction in *PLP1* expression on single EGC transcriptomes from the report by Kinchen *et al* [29], and from the integrated scIBD dataset by Nie *et al* (Supplemental Fig. 2D-E) [20]. In addition, we also reanalysed 13,440 single EGC transcriptomes from in the dataset by Thomas *et al* [30], confirming the significant upregulation of *CXCL9* and cell death pathway signature expression and a downregulation of *PLP1* in EGC from inflamed UC and CD patients (Supplemental Fig. 2F-H).

Taken together, our data identify a preferential IBD type-related induction of specific EGC markers along with an induction in cell death pathway expression specifically in activated EGC from inflamed IBD (UC > CD) patients.

### Rapid induction of EGC activation and cell death in response to Th1/Th17-driven small intestinal inflammation

To gain mechanistic insights, we screened several mouse models of experimental intestinal inflammation for canonical EGC markers. For this, we choose different preclinical modelling paradigms of intestinal inflammation to investigate the impact on EGC homeostasis. These included the Th1-driven ileitis model of *E. vermiformis* infection (E.veri) [31], the TNF dominant ileitis model of constitutive TNF production (TNFdARE) [32], the Th1/Th17-driven model of acute ileitis via *in vivo* T-cell activation (aCD3) [33], and the IL-33 dominant model of ablating *Casp8* specifically in intestinal epithelial cells (Casp8_Ile) [34]. We recently reported analysis of the whole transcriptomes from these models leveraging which, we performed the initial screening [35]. This analysis showed a rapid (within 6h) and strong increase in the aCD3 model for the mouse EGC activation marker *Gfap*, and a repression of *Plp1*, akin to our observations from patient datasets. The infectious model E.veri also paralleled the strong repression for certain EGC markers but showed only mild induction of the activation marker *Gfap.* Compared with these, the Casp8_Ile had a mild impact on the overall expression of the EGC markers (Fig. 3A). Similar to our earlier report, analysis of several widely used colitis models showed a distinct impact on EGC markers with a hallmark *Gfap* induction and *Kcna1* repression only observed in the trinitrobenzene sulfonic acid (TNBS)-induced colitis (Supplemental Fig. 3A) [35, 36].

**Figure 3.**
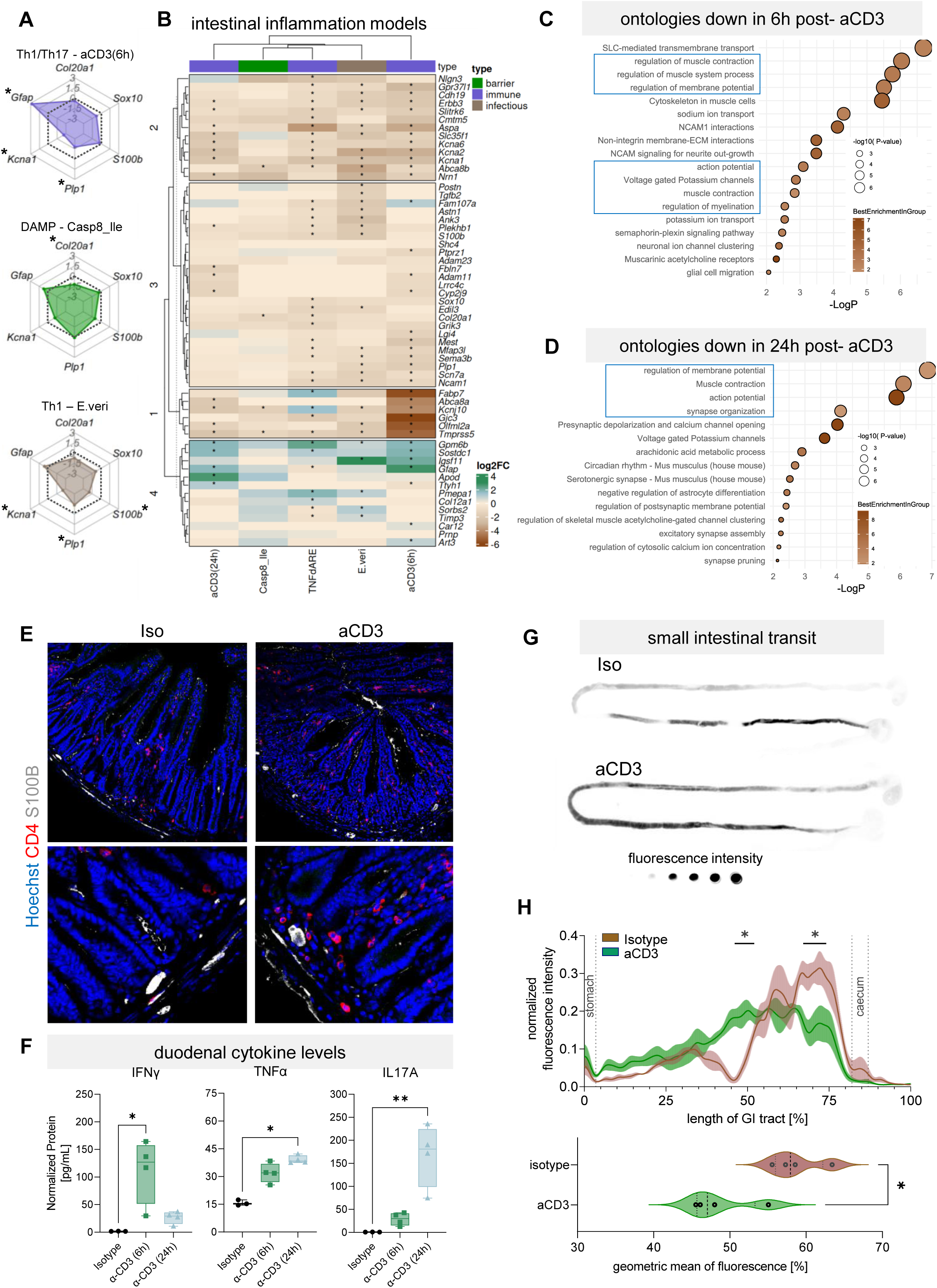
Rapid impact on enteric glial cell (EGC) homeostasis and gut motility by Th1/Th17-driven intestinal inflammation. **A)** Radar charts showing log2 fold changes of key mouse EGC genes in different modelling paradigms of intestinal inflammation [aCD3 = *in vivo* anti-CD3 antibody-induced T-cell driven intestinal inflammation, Casp8_Ile = ileitis induced by intestinal epithelial specific-ablation of *Casp8*, E.veri = *Eimeria vermiformis* infection-induced intestinal inflammation] [35]. Asterisk indicate p-values < 0.05 from the DESeq2 Wald statistic. **B)** Heatmap showing log2 fold changes of mouse EGC-enriched genes extracted from sorted EGC transcriptomes published previously [7] and tracked on indicated modelling paradigms of intestinal inflammation [aCD3, Casp8_Ile, E.veri as in A above. 6h and 24h indicate 2 different time points post aCD3 injection; TNFdARE = intestinal inflammation induced by constitutive TNF expression [32]. Asterisk indicate p-values < 0.05 from the DESeq2 Wald statistic. **C-D)** Selected gene ontologies downregulated in the transcriptomes at the 6h (C) and 24h (D) time points after aCD3-induced gut inflammation. Blue boxes indicate significantly enriched ontologies of interest. **E)** Representative photomicrographs of immunostained jejunal tissues from the aCD3 versus isotype (Iso) control tissues after 6h, showing high levels of CD4+ positive cells in close proximity to S100B positive plexi compared with isotype treated controls. **F)** Levels of indicated cytokines measured from duodenum of mice treated with Iso versus aCD3 groups from the indicated time points. Data are derived from three to four biological replicates per group. Asterisk indicate p < 0.05 from one-way ANOVA followed by the Kruskal-Wallis post-hoc test. **G & H)** Assessment of small intestinal transit in Iso and aCD3 groups (G) representative images of dye transit in the gut tissues of indicated groups, and (H) normalized intensity along the gut (top) with geometric means of fluorescence (bottom). Asterisk indicate p < 0.05 between the groups from the two-way ANOVA followed with Tuckey’s post-hoc test (top) and the non-parametric Mann Whitney test (bottom) from four independent biological replicates per group.

To broaden the available EGC activation markers in preclinical models, we adopted the same approach as that used for analysing the human cohorts. For this, we extracted top EGC enriched genes from sorted EGC versus non-EGC in the setting of preclinical gut inflammation, reanalysed from Progatzky *et al* (Supplemental Fig. 3B) [7]. This analysis showed a rapid regulation in several specific EGC markers, some of which were conserved with the changes seen in the IBD patient datasets. Intrigued by its rapidity and to minimize the influence of the environmental and microbial components we focused on the aCD3 model to directly assess the impact of Th1/Th17-driven gut inflammation on EGC for further characterization. These included the induction of *Gfap,* and *Art3* and a downregulation of *Plp1* and *Lgi4* among others (Fig. 3B). Selected pathway ontologies from the transcriptomes of the 6h and 24h aCD3 gut transcriptomes showed an overall repression in pathways associated with neuromuscular function and myelination (Fig. 3C, D). These transcriptomic changes were accompanied by CD4⁺ T-cell infiltration in the vicinity of enteric plexuses, increased levels of cytokines IFN-γ, TNF, and IL-17A in the tissues, and muscularis inflammation at 6h and 24h after aCD3 induction (Fig. 3E-F, Supplemental Fig. 3C-E). Moreover, the rapid repression in *Plp1* transcripts shown earlier was accompanied with a reduction in PLP1 protein level (Supplemental Fig. 3F). Alongside these EGC related transcriptomic, cellular, and protein level alterations, we also observed a rapid and significant reduction in small intestinal motility in the aCD3 model, a crucial process regulated by EGC (Fig. 3G, H).

Given these rapid alterations in EGC homeostasis and the functional consequences in response to elevated Th1/Th17 activation, we established the aCD3 model in *Plp1*Cre*ERT*; Rosa26-tdTomato+ reporter mice enabling us to sort tdTomato+ EGCs for analysis (Fig. 4A, Supplemental Fig. 4A). Six hours post-aCD3, the sorted EGC showed a significant alteration in their transcriptomes compared to isotype (Iso) controls with the first principal component accounting for 81% of the variation among the groups (Fig. 4B). As expected from tissue cytokine measurements, several interferon-driven genes such as *Cxcl9* and *Cxcl10,* the EGC activation marker *Gfap* (7.7 log2 fold), and interestingly the necroptotic executioner *Mlkl* (4.4 log2 fold) were highly upregulated (Fig. 4C). Consequently, gene ontology analysis showed the upregulation of interferon and TNF signalling, gliogenesis, and necroptotic processes (Fig.4D).

**Figure 4.**
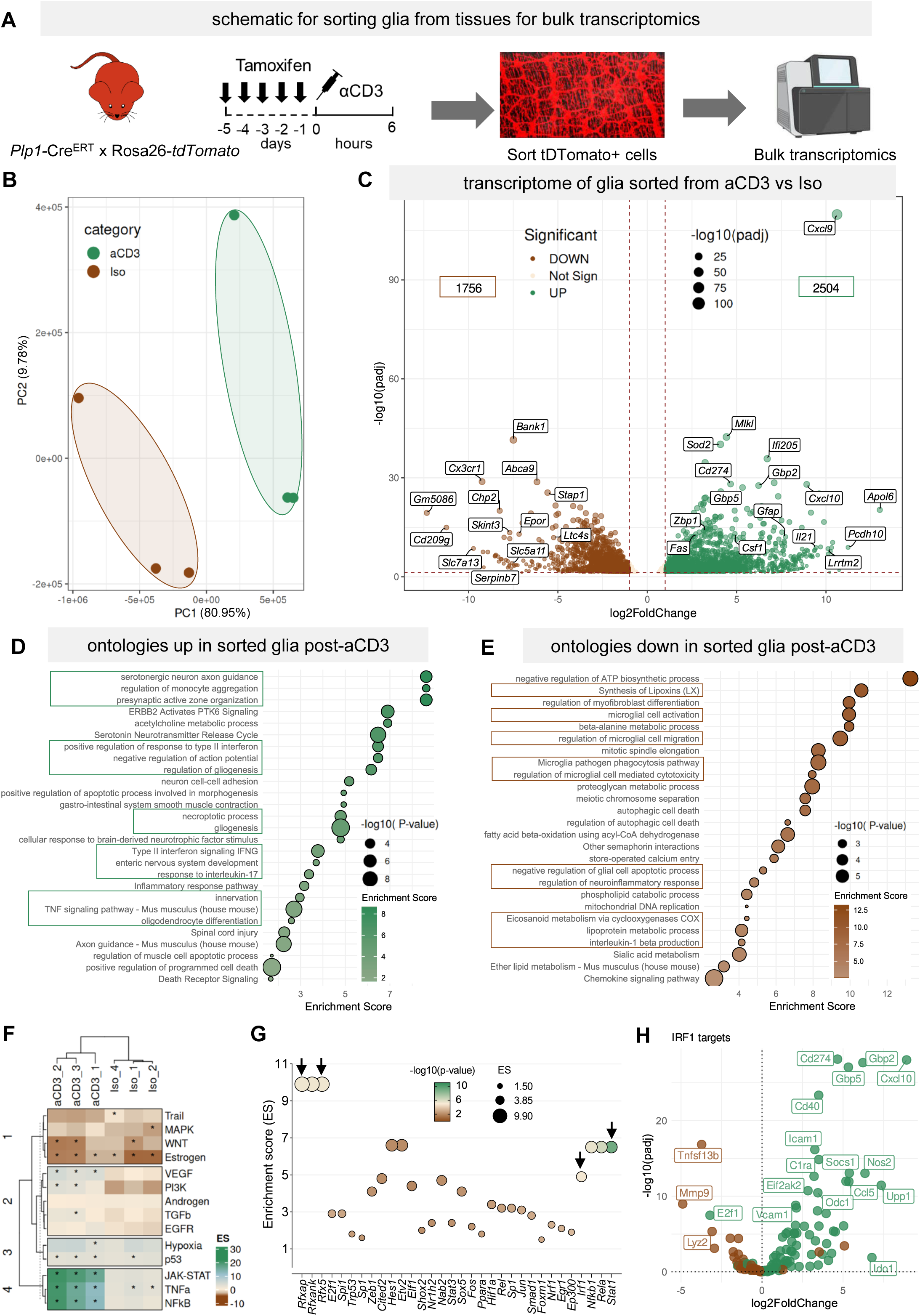
Sorted enteric glial cell (EGC) transcriptomes reveal immune-driven activation and necroptosis programs. **A)** Schematic depiction of fluorescence activated cell sorting of EGC from *Plp1CreERT*; tdTomato+ glia reporter mice in the aCD3 model for bulk transcriptomics. **B)** The first and second principal components (PC) from the transcriptomes of the isotype (Iso) and the aCD3 treated mice. **C)** Volcano plot showing transcriptomic impact of aCD3 on sorted EGC versus Iso control EGC. Data is derived from three independent biological replicates per group. Dashed brown line along the x-axis denotes adjusted p-value threshold of 0.05 from the DESeq2 Wald statistic. Vertical dashed brown lines indicate absolute log2 fold change threshold of 0.58. **D & E)** Selected gene ontologies that are (D) upregulated and (E) downregulated in the sorted EGC from aCD3 versus Iso groups. Boxes with green and brown outlines indicate significantly enriched ontology terms of interest. **F)** Heatmap showing enrichment scores (ES) from the inference of upstream pathways contributing to the observed transcriptomic changes in (C) above, via PROGENy [37]. Asterisk indicate p < 0.05 from the multivariate linear modelling. **G-H)** Analysis of inferred transcription factors (TFs) contributing to the transcriptomic changes in (C) above, via CollectTri [39] (G) Dot plot showing enrichment scores (ES) and p-values for indicated TFs and (H) Volcano plot for predicted IRF1 targets (positive targets = green dots and negative targets = brown dots) and their log2 fold changes from the transcriptomes of EGC sorted from aCD3 versus Iso treated mice as shown in (C) above.

Interestingly, several downregulated genes such as *Chp2*, *Cx3cr1, Stap1* and *Epor* hinted at repressed adaptive immune and stress responses (Fig. 4C). Similar to the opposite regulation of *Gfap* and *Cx3cr1* on our sorted EGC transcriptomes, reanalysis of a previous scRNA-Seq study also showed differential EGC-activation-related expression of these, and other immune homeostatic and metabotropic genes (Supplemental Fig. 4B [7]). Altogether, several downregulated ontologies related with microglia activation and regulation of neuroinflammatory and stress response genes (Fig. 4E). Pathway and transcription factor activity inference analysis from the transcriptomes of our sorted EGC reconfirmed TNF, JAK-STAT, and NFkB as upstream pathways and STAT1, IRF1, and the RFX family as the main transcription factors (TF) contributing to the transcriptomic alterations seen in the sorted EGC from the aCD3 model (Fig. 4F-H) [37, 38, 39, 40]. A high enrichment score was also observed for the TF HES1, which has known roles in the regulation of ENS development and EGC differentiation (Fig. 4G).

These findings pointed at an induction in pathways related with EGC activation and necroptosis upon Th1/Th17-induced intestinal inflammation. We could also confirm such EGC activation-associated elevation in cell death pathway signature by reanalysing the dataset by Progatzky *et al*. [7] (Supplemental table ST2). Interestingly, the cluster with elevated cell death from this analysis also showed increased expression of *Ifngr1* and *Tnfrsf1a* receptors, which was in line with our observations from our sorted EGC transcriptomes from the aCD3 model (Fig. 5A, 4C-D).

**Figure 5.**
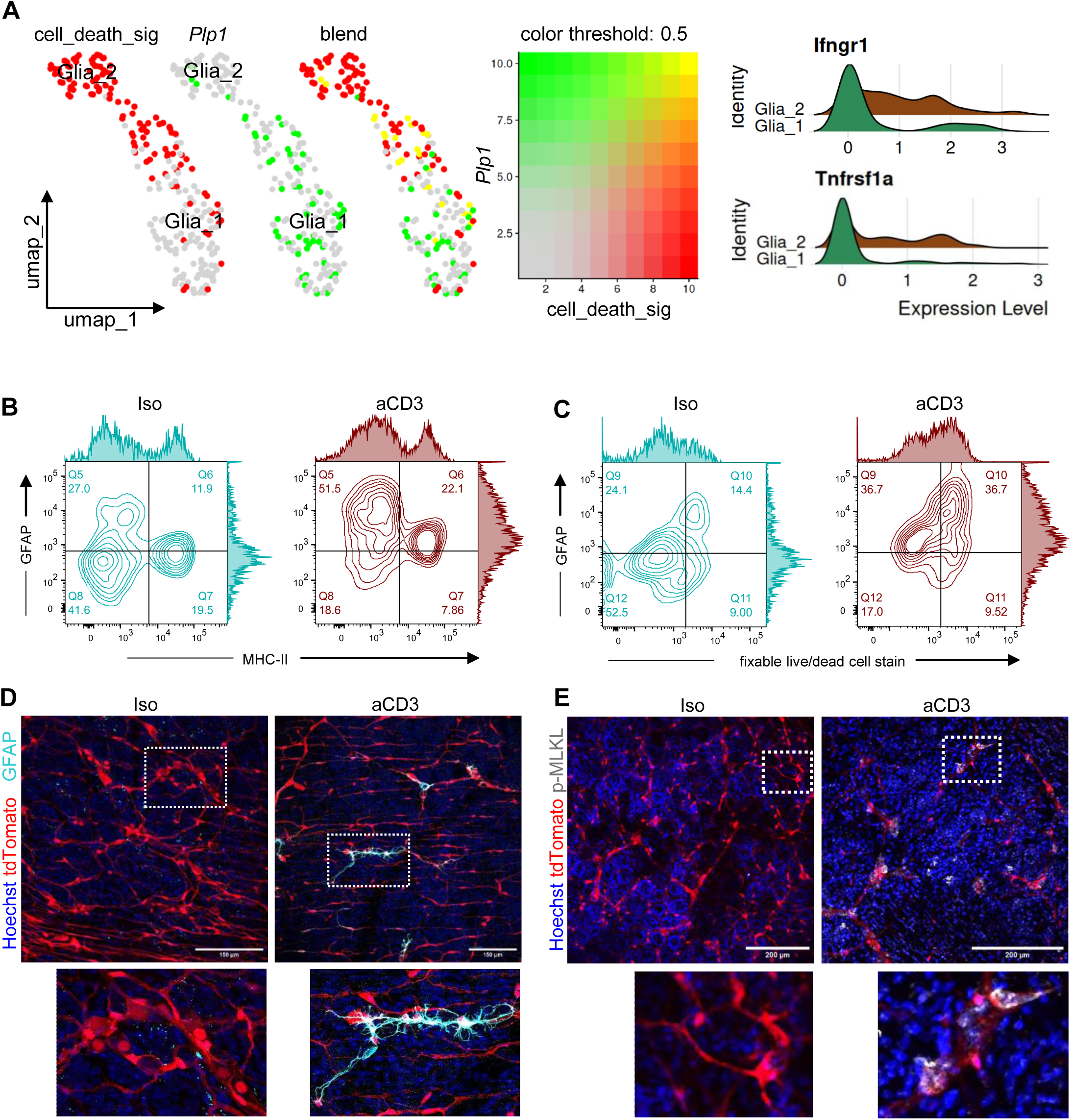
Induction of necroptosis in activated enteric glial cells (EGC) in Th1/Th17-driven intestinal inflammation. **A)** Uniform manifold approximation and projection (umap) plots from single mouse EGC transcriptomes from Progatzky *et al*. showing the thresholded expression levels for the mean cell death signature and *Plp1* alongside ridge plots for *Ifngr1* and *Tnfrsf1a* showing cluster of activated EGC (Glia_2) [7] with higher expression of the cell death pathway signature and lower *Plp1* expression. **B-C)** Density plots from flow cytometry analyses of indicated EGC activation markers and fixable cell death dye gated for tdTomato+ and isolated from the muscularis of mice 6h post-injection of anti-CD3 antibody (aCD3) versus isotype (Iso) control. **D-E)** Photomicrographs from whole-mount immunofluorescence staining showing colocalization of the tdTomato EGC reporter with mouse EGC activation marker GFAP (D), and (E) that of the necroptosis executioner phospho-Ser345 MLKL from the submucosal plexi of aCD3 and Iso controls.

Next, we directly measured whether such activation-associated EGC death was indeed observable in the acute Th1/Th17-driven intestinal inflammation. In mouse EGC, activation is known to be associated with elevation in GFAP and MHC-II [6]. Hence, flow cytometry analysis was performed on tdTomato+ EGC for activation coupled with fixable cell death dye from gut tissues of aCD3 versus Iso-challenged *Plp1*^Cre*ERT+*^; Rosa26^tdTomato+^ reporter mice. Analysis revealed a two-fold induction in tdTomato+, GFAP+, MHC-II+ EGCs and a 2.5-fold increase in tdTomato+, GFAP+, fixable dead stain+ EGC in the aCD3 versus the Iso group (Fig. 5B, C, Supplemental Fig. 5 A-B). Moreover, whole mount immunostaining showed higher GFAP+ tdTomato+ EGC in the submucosal plexuses of aCD3, but not Iso treated mice (Fig. 5D). An elevated immunopositivity for phosphorylated Ser345 MLKL (pMLKL) was seen in tdTomato+ cells in the aCD3, but not in the Iso group (Fig. 5E). This was accompanied by a generalized reduction in the overall tdTomato signal in the aCD3 compared with Iso tissues (Fig. 5D, E). These data indicate acute cytokine-mediated EGC activation in conjunction with EGC regulated necrosis in a Th1/Th17 dominant intestinal inflammation.

### IFN-γ and TNF drive activation and necroptosis induction in sorted EGC *ex vivo*

To directly evaluate the impact of Th1/Th17 cytokines on the EGC activation and cell death, we established protocols to culture and analyse FACS-sorted EGC from the muscularis of *Plp1*^Cre*ERT*+^; Rosa26^tdTomato+^ reporter mice (Supplemental Fig. 6A). The sorted tdTomato+ cells were confirmed immunopositive for the canonical EGC markers SOX10 and S100B with some TUBB3 expression after 10 days in culture (Supplemental Fig. 6B-E) [41]. *Ex vivo* culturing enabled the direct investigation of how IBD-associated cytokines impact EGC. Based on our data so far and on previous reports, we selected IFN-γ, TNF, IL-17A, and IL1-β [7, 24, 42, 43]. Sorted EGCs in *ex vivo* culture displayed strong transcriptomic responses to IFN-γ, TNF, and IL1-β, whereas IL-17A elicited only a weak transcriptomic change (Fig. 6A-B, Supplemental Fig. 6F, G). Both IFN-γ and TNF led to common upregulation of inflammation associated transcripts including chemokines *Cxcl9* and *Ccl2*, similar to that found our earlier finding from sn– and sc-RNA-Seq datasets from inflamed human EGC (Supplemental Fig. 6H). There was also a very high degree of transcriptomic overlap (R^2^ 0.927) between the IL1-β and TNF stimulated EGC owing to autocrine TNF expression upon IL1-β stimulation (Supplemental Fig. 6I). Given these data and the simultaneous presence of both IFN-γ and TNF at high levels in the tissues of the aCD3 model (Fig. 3F), we postulated a combinatorial action of these cytokines in shaping EGC responses during inflammation. To investigate this, we analysed transcriptomes of sorted EGC upon IFN-γ and TNF co-stimulation. Interestingly, co-stimulation with IFN-γ and TNF led to a strong upregulation in several genes including *Mlkl*, *Cd274*, *Gbp2*, and *Ifi205* that were also highly upregulated in the transcriptomes of EGC sorted from the aCD3 model (Fig. 6C and Fig. 4C). Surprisingly, IFN-γ and TNF co-stimulation caused several gene ontologies related with necroptotic cell death to be upregulated. Most interestingly, the gene ontology terms for inhibition of Caspase-8 activity, necroptotic signalling, and programmed necrotic cell death were upregulated besides those indicative of EGC activation such as MHC Class II protein complex assembly (Supplemental Fig. 6J). This was interesting given that inhibition of Caspase-8 activity promotes necroptosis [44]. Moreover, the IFN-γ and TNF co-stimulation also led to a strong downregulation of several key genes *Ret*, *Phox2b*, *Gpr37l1*, *Edn3*, and *Ascl1* involved in controlling ENS homeostasis (Fig. 6C). This was reflected in significant enrichment in selected downregulated gene ontology terms related with glial cell and enteric nervous system differentiation (Supplemental Fig. 6K).

**Figure 6.**
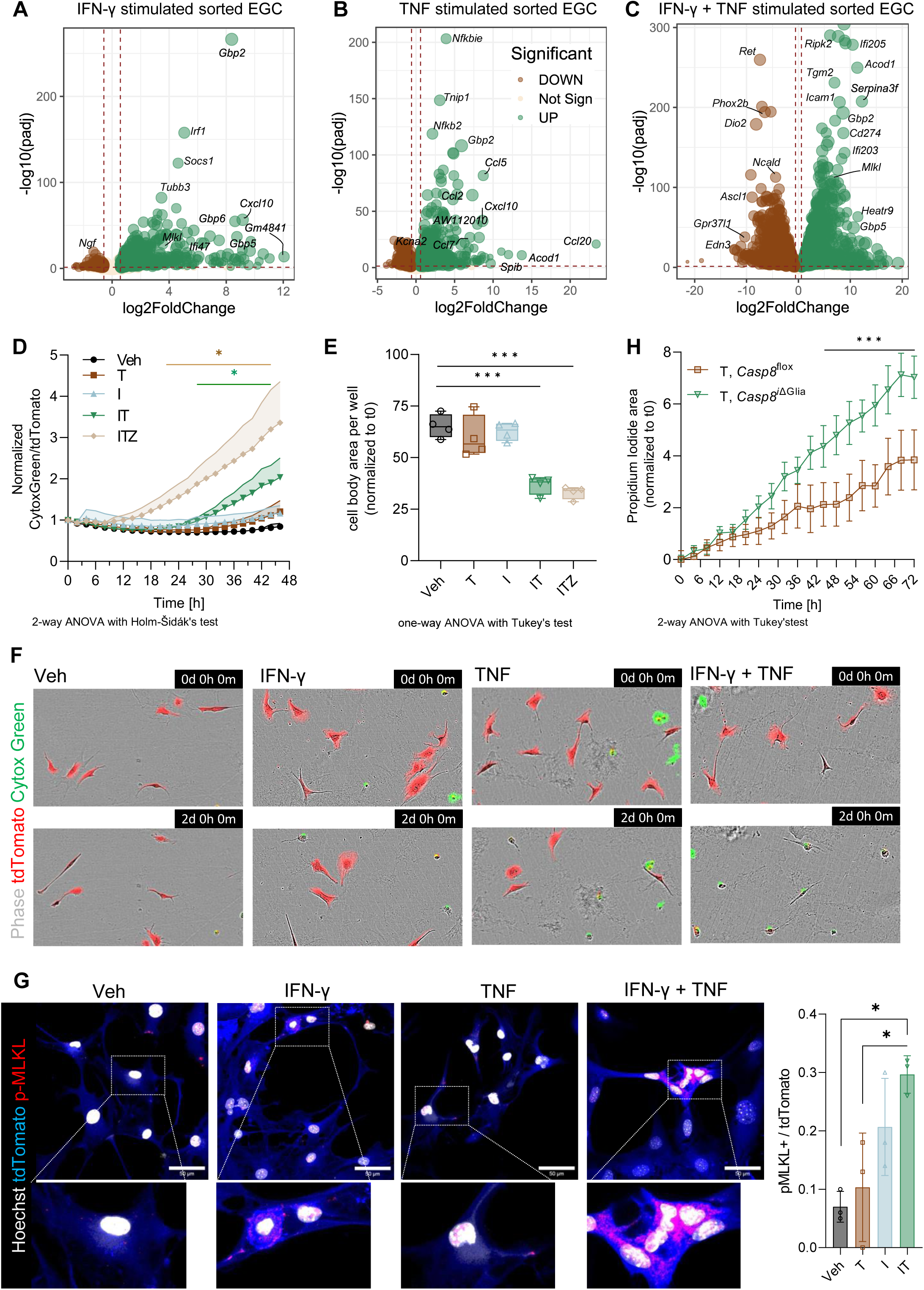
IFN-γ and TNF coordinate caspase-independent death of enteric glial cells (EGC) accentuated by glial *Casp8* ablation. **A-C)** Volcano plots showing transcriptomic impact of the cytokines (A) IFN-γ, (B) TNF, and their combination (C) IFN-γ + TNF on sorted EGC *ex vivo*. Data is derived from three to four independent biological replicates per group, dashed brown line along the x-axis denotes adjusted p-value threshold of 0.05 from the DESeq2 Wald statistic. Vertical dashed brown lines indicate absolute log2 fold change threshold of 0.58. **D)** Live cell tracking of cell death in sorted tdTomato+ EGC stimulated with recombinant mouse cytokines TNF (T), IFN-γ (I) and the pan caspase inhibitor zVAD-fmk (Z), either alone or in combinations as indicated, over a period of 48 hours. **E)** Analysis of cell body area at the end of the experiment in (D), normalized to day 0. For (D) and (E) n = 4 biological replicates per condition, information on statistical testing is included below the figures. **F)** Representative images from indicated time points and treatment groups of the sorted EGC live cell imaging experiment in (D) and (E) [Veh = vehicle control]. **G)** Representative photomicrographs of tdTomato+ sorted EGC, stimulated *ex vivo* with the indicated cytokines or vehicle (Veh) control for 48 hours and immunostained for phosphorylated Ser345 MLKL (pMLKL) followed by quantification of the pMLKL to tdTomato ratio in each group for three independent biological replicates per treatment group. Asterisk indicate p < 0.05 from one way ANOVA followed by Tuckey’s post-hoc test. **H)** Live cell monitoring for cell death via propidium iodide in plexanoids isolated from littermate mice either with glia-specific inducible ablation of *Casp8* (*Plp1^CreERT+^*; *Casp8*^fl/fl^ = *Casp8^i^*^ΔGlia^) or Cre controls (*Plp1^CreERT-^*; *Casp8*^fl/fl^ = *Casp8*^flox^). Both groups were treated with 4-hydroxytamoxifen (500nM) before stimulation with recombinant mouse TNF (T) (50ng.ml^-1^). Data is from three independent biological replicates per group. Information on statistical testing is included below the figure.

These data confirmed that combinatorial action of IFN-γ and TNF drives EGC activation, necroptosis induction, and reduction in glial replenishment during intestinal inflammation. Next, we directly assessed cell viability via time lapse imaging of sorted tdTomato+ EGC over a 48h period in response to IFN-γ, TNF, or their combination (Fig. 6D-F). Analysis revealed that neither TNF nor IFN-γ on their own were enough to drive actual cell death of EGC. However, when EGC were treated with a combination of TNF and IFN-γ, a significant increase in cell death and a consequent reduction in the area covered by EGC were observed (Fig. 6D-F). Attempts to rescue this TNF and IFN-γ induced cell death from apoptosis or pyroptosis via pharmacological blockade of all caspases using zVAD-fmk (Z) failed (Fig. 6D-E). In fact, the addition of the pan-caspase inhibitor led to a further accentuation of cell death, a characteristic of regulated necrosis (Fig. 6D-E) [45]. Based on these data from the IFN-γ and TNF co-stimulated EGC showing elevated *Mlkl* expression and the ontology term inhibition of Caspase 8 activity, we analysed the functional induction of MLKL by immunostaining for pMLKL. A significant elevation in pMLKL, and pMLKL to tdTomato signal intensity was detectable only in sorted EGC co-stimulated with IFN-γ and TNF (Fig. 6G).

Given that the coordinated action of IFN-γ and TNF showed transcriptomic induction of the ontology term inhibition of Caspase 8 along with functional induction of necroptosis, we hypothesized that an ablation of *Casp8* in EGC would potentiate this susceptibility. In order to investigate this, we generated *Plp1*^Cre*ERT+*^*; Casp8^flox/flox^* (*Casp8^i^*^ΔGlia^) mice, in which *Casp8* deletion can be induced specifically in glial cells. Caspase-8 is a critical regulator of extrinsic cell death downstream of TNF and reports from others and us have shown that *Casp8* ablation predisposes to necroptosis in various cell types [34, 44, 46]. Time lapse tracking of cell death from plexanoid cultures (Supplemental Fig.6L, M) derived from the muscularis of *Casp8^i^*^ΔGlia^ and *Casp8*^flox^ mice revealed that TNF alone was sufficient to induce significant elevation in cell death without the additional requirement of IFN-γ (Fig. 6H). As expected, given that EGC necroptosis was already elevated in in the aCD model, the aCD3-induced muscularis inflammation, disease scores and muscle thickness remained comparable between *Casp8^i^*^ΔGlia^ and *Casp8^flox^* mice. However, a higher presence of red blood cells in the lumen and tissue parenchyma was observed in the *Casp8^i^*^ΔGlia^ group (H&E –bleeding) (Supplemental Fig.6N-P). Taken together, these data indicate that pathological programmed necrosis of EGC in the gut is coordinated by IFN-γ and TNF and that *Casp8* ablation renders EGC susceptible to TNF-induced cell death without the requirement of IFN-γ.

Overall, our data identify new EGC-specific activation markers across clinical and preclinical IBD and identify regulated necrosis of activated EGC coordinated by IFN-γ and TNF in acute intestinal inflammation.

## Discussion

The precise roles and the impact of inflammation on EGC in IBD are underrecognized. Traditionally considered neuroprotective and neurotrophic, the immune and epithelial regulatory roles of activated EGC in inflamed guts are just beginning to be uncovered [47]. Enteric and peripheral neuropathy is frequently observed in IBD, although the exact origins remain elusive with speculation that the path to immune attack on neuroglia might be paved first via dysfunctional immune-EGC crosstalk. There is growing evidence from mouse models suggesting that specifically, the *Gfap* expressing, activated EGC exert several beneficial functions. This includes activated EGC-derived Wnt in improved epithelial recuperation post-injury as well as improved helminth clearance [3, 48]. Moreover, in mice activated EGC can induce tolerogenic T-cells via MHC-II mediated intrinsic antigen presentation [6]. Despite these reports, recent single cell studies have cast some doubt on the specificity of GFAP in labelling EGC [8, 9, 49]. Therefore, there is an emergent need for the identification of EGC-specific markers which are also inflammation associated. Our study addressed this need by extracting EGC enriched genes from an integrated collection of IBD single cell transcriptomes. This led us to some interesting an unexpected EGC-enriched genes with as yet unknown implications in immune and epithelial crosstalk. For instance, we found that EGC from inflamed IBD samples displayed higher expression of *SPP1* on multiple datasets from colon tissues of IBD. Previous reports indicate that in mice small intestinal intra-epithelial lymphocytes can express *SPP1*. However, its dominant cellular source in the human colon has not been defined. Similarly, *WNT6*, which was induced in EGC from inflamed IBD samples, is a crucial epithelial morphogen, which in the small intestine is sourced from Paneth cells, however its source in the colon is unknown.

Another important function of EGC, which has become increasingly appreciated in recent years is their ability to carry out injury-induced neurogenesis, replenishing lost enteric nervous system (ENS) cells in mouse models [50, 51, 52, 53]. This is enabled due to the existence of a multipotent proteolipid protein 1 (PLP1) expressing glia-like quiescent cell in the myenteric plexus. Data from IBD patients has so far not reported the presence of such a population. One of the EGC subclusters identified on our snRNA-Seq is a *PLP1* high cluster, which simultaneously co-expressed a number of neural crest precursor genes including *PHOX2B*, *SOX2*, and *RET*. Moreover, this cluster was rich in patient EGC compared with those from non-IBD controls indicating that inflammation may trigger the emergence of neurogenic EGC in IBD. Ours is the first report to describe such a cluster, presumably given that we analysed single nuclei from full-thickness gut tissues, avoiding the cell health related losses that affect single cell isolation protocols. Interestingly, this cluster was spared from the induction in cell death pathway signature that was observed in the activated EGC clusters.

Pathological, caspase-independent TNF– and IFN-γ-driven cell death has been previously reported in brain oligodendrocytes [54]. However, ours is the first report on extrinsic necroptosis induction in response to inflammatory cytokines in EGC. This is particularly relevant in IBD, where immune cell infiltration into ENS plexuses alters neuroglial function via local cytokine production [55], but its impact on EGC health has been largely ignored. Elevation of TNF and IFN-γ in tissues is known to synergistically coordinate inflammatory, necroptosis via heavily inducing expression and phosphorylation of the necroptosis executioner MLKL [56]. Both TNF and IFN-γ levels were elevated in the tissues of our Th1/Th17 intestinal inflammation model concomitant with activation of EGCs, increased transcription, and phosphorylation on of MLKL. Other gastrointestinal cell types have also been shown to undergo necroptosis in response to interferon-induced MLKL upregulation, and our findings extend this paradigm to EGCs [46, 57, 58]. Overall, our study provides new insights into the inflammatory regulation of EGC turnover via specific culling of activated EGC in Th1-Th17 dominant intestinal inflammation.

A differential impact of inflammation on EGC subtypes and the functional consequences to gut physiology in IBD remains an open question. It is likely that EGC turnover is influenced in patients who respond to anti-TNF therapy via reduced activation and associated cell death. Given that EGC subsets regulate gut motility, it is interesting to note that previous and ongoing clinical trials in CD have proposed that magnetic resonance imaging-assisted monitoring of gastrointestinal motility might be a predictor of anti-TNF therapy response [59, 60]. The protection from cytokine-induced EGC death may extend to EGC clusters with different functions. Further studies aimed at understanding the molecular and functional diversity of EGC subtypes and their implications in IBD will provide an improved understanding of the complex crosstalk between the enteric nervous system’s role in maintaining neuro-immune-epithelial homeostasis in the gut.

## Materials and methods

Detailed materials and protocols are provided in the supplemental information section.

## Data availability

All of the raw sequencing data from mouse sorted EGC from the aCD3 model and sorted-cultured EGC stimulated with cytokines that were generated from this are accessible via ArrayExpress (https://www.ebi.ac.uk/biostudies/arrayexpress) at the following accession numbers (E-MTAB-14876, E-MTAB-14879). Raw sequence data from mouse colitis and enteritis models analysed in this study are also available via ArrayExpress (AcDSS, cDSS, and TC: E-MTAB-14306; Everm: E-MTAB-14297; Hhepa: E-MTAB–14316; OxC, Crode: E-MTAB– 14312; Casp8(Col) and Casp8_Ile: E-MTAB-14318; AcTNBS, cTNBS: E-MTAB-14329; *Tnf*^ΔARE^Col and *Tnf*^ΔARE^Ile: E-MTAB-14325, anti-CD3: E-MTAB-14831). Publicly available mouse sorted EGC bulk RNA Seq datasets analysed in this study can be accessed via the gene expression omnibus (GEO) https://www.ncbi.nlm.nih.gov/geo/ with the accession number GSE182708. Patient microarray and bulk RNA Seq data for publicly available IBD cohorts analysed in this study can be accessed via GEO (Microarrays: GSE10191, GSE10616, GSE6731, GSE4183, GSE6731, GSE9686; Bulk RNA Seq: GSE193677). Transcriptomic data from the IBDome cohort (v1.0.0) are available at https://ibdome.org. Publicly available processed human single cell RNA sequencing datasets and integrated scRNA Seq datasets used in this study can be accessed using the following accession numbers or weblinks: Kinchen *et al.*: GSE114374 [29], Nie *et al.*: http://scibd.cn/ [20], Thomas *et al.*: https://doi.org/10.5281/zenodo.13768607 [30], Oliver *et al.*: https://gutcellatlas.org/pangi.html [28]. Access to the snRNA-Seq data generated in this study are available upon request.

## Code availability

Code used to generate the analyses in this study is accessible on GitHub via the following links https://github.com/jaypaty/EGC_paper, https://github.com/jaypaty/microarray_IBD_EGC_2025

## Acknowledgements

We acknowledge the support provided for animal husbandry and rearing by Eva Wellein and Melanie Ziegler. We also acknowledge the work and support of Uwe Appelt, Markus Mroz, and Samaneh Sotoodeh Sheyjani of the core unit for cell sorting and immune monitoring of the Friedrich-Alexander-Universität Erlangen-Nürnberg (FAU). Moreover, we thank Fabrizio Mascia for his outstanding technical assistance and support in this project. We thank the MDC/BIH Genomics Platform, Berlin (FacilityID=1565, The CoreMarketplace: MDC & BIH Technology Platform Genomics) for technical support and sequencing.

## Funding support

This work was funded by the Deutsche Forschungsgemeinschaft (DFG, German Research Foundation) – TRR241 375876048 (A03, A05, A08, B02, B05, Z03), SFB1181 (C05). The project was further supported by 505539112 (KFO5024 Project – A3). A.N.H is supported by a Lichtenberg fellowship and “Corona Crisis and Beyond” grant by the Volkswagen Foundation, a BIH Clinician Scientist grant and INST 335/597-1. FP is funded by Wellcome (226579/Z/22/Z), Kennedy Trust for Rheumatology Research (KENN 21 22 11) and Versus Arthritis (22981). Moreover, the project was supported by the DFG grant covering sequencing costs to B.S and C.B Projektnummer 418055832. The project was also supported by the Interdisciplinary Centre for Clinical Research (IZKF: A76, A93, J96, and ELAN P120). The authors declare no financial and non-financial conflicts of interest arising from this work.

## Author Contributions

The study was conceived by MB and JVP and planned by MB, HL, KAP, and the IBDome Consortium. Data was acquired by MB, HL, KAP, MGA, CP, LE, RGB. Data was analysed by MB, KAP, MGA, CP, and JVP. Data interpretation and help with protocols was provided by IBDome Consortium, MB, HL, KAP, MGA, PT, MJ, MR, AM, FP, SK, RGB, LE, CB, CR, LSL, CG, KH, ZT, ANH, JVP. The figures were prepared by MB, HL and JVP. MB and JVP wrote the manuscript. Help with manuscript editing and critical review was provided by RGB, LE, AAK, ANH, MR, AM, FP, SK, BS, CB, CSNK, DCL, and MFN. Help with securing funds, study supervision, and key resources JVP, CSNK, CB, BS, ANH, RA, DCL, CG, CR, LSL, KH, ZT, BS, and MFN, and the IBDome Consortium.

## Consortia associated with this study

The TRR241 IBDome Consortium (in alphabetical order):

Imke Atreya1,2, Raja Atreya1,2, Petra Bacher3,4, Christoph Becker1,2, Christian Bojarski5, Nathalie Britzen-Laurent6, Caroline Voskens7, Hyun-Dong Chang8,9, Andreas Diefenbach10, Claudia Günther1,2, Ahmed N. Hegazy5, Kai Hildner1,2, Christoph S. N. Klose10, Kristina Koop1,2, Susanne M. Krug11, Anja A. Kühl12, Moritz Leppkes1,2, Rocío López-Posadas1,2, Leif S. Ludwig13,14, Clemens Neufert1,2, Markus Neurath1,2, Jay V. Patankar1,2, Christina Plattner15, Magdalena S. Prüß5, Andreas Radbruch8, Chiara Romagnani16, Francesca Ronchi10, Ashley D. Sanders5,13,14, Alexander Scheffold4, Jörg-Dieter Schulzke11, Michael Schumann5, Sebastian Schürmann17, Britta Siegmund5, Michael Stürzl6, Zlatko Trajanoski15, Antigoni Triantafyllopoulou8,10,18, Maximilian Waldner1,2, Carl Weidinger5, Stefan Wirtz1,2, Sebastian Zundler1,2

1. Department of Medicine 1, Universitätsklinikum, Friedrich-Alexander-Universität Erlangen-Nürnberg, Erlangen, Germany
2. Deutsches Zentrum Immuntherapie, Universitätsklinikum, Friedrich-Alexander-Universität Erlangen-Nürnberg, Erlangen, Germany
3. Institute of Clinical Molecular Biology, Christian-Albrecht University of Kiel, Kiel, Germany.
4. Institute of Immunology, Christian-Albrecht University of Kiel and UKSH Schleswig-Holstein, Kiel, Germany.
5. Charité – Universitätsmedizin Berlin, corporate member of Freie Universität Berlin and Humboldt-Universität zu Berlin, Department of Gastroenterology, Infectious Diseases and Rheumatology, Berlin, Germany
6. Department of Surgery, Universitätsklinikum, Friedrich-Alexander-Universität Erlangen-Nürnberg, Erlangen, Germany
7. Department of Dermatology, Universitätsklinikum, Friedrich-Alexander-Universität Erlangen-Nürnberg, Erlangen, Germany
8. Deutsches Rheuma-Forschungszentrum, ein Institut der Leibniz-Gemeinschaft, Berlin, Germany
9. Institute of Biotechnology, Technische Universität Berlin, Germany
10. Charité – Universitätsmedizin Berlin, corporate member of Freie Universität Berlin and Humboldt-Universität zu Berlin, Institute of Microbiology, Infectious Diseases and Immunology
11. Charité – Universitätsmedizin Berlin, corporate member of Freie Universität Berlin and Humboldt-Universität zu Berlin, Clinical Physiology/Nutritional Medicine, Department of Gastroenterology, Rheumatology and Infectious Diseases, Berlin, Germany
12. Charité – Universitätsmedizin Berlin, corporate member of Freie Universität Berlin and Humboldt-Universität zu Berlin, iPATH.Berlin, Berlin, Germany
13. Berlin Institute of Health at Charité – Universitätsmedizin Berlin
14. Max Delbrück Center für Molekulare Medizin, Berlin
15. Biocenter, Institute of Bioinformatics, Medical University of Innsbruck, Innsbruck, Austria
16. Charité – Universitätsmedizin Berlin, corporate member of Freie Universität Berlin and Humboldt-Universität zu Berlin, Institute for Medical Immunology
17. Institute of Medical Biotechnology, Friedrich-Alexander-Universität Erlangen-Nürnberg, Erlangen, Germany
18. Charité – Universitätsmedizin Berlin, corporate member of Freie Universität Berlin and Humboldt-Universität zu Berlin, Department of Rheumatology and clinical Immunology

## Notes

### Competing Interest Statement

The authors have declared no competing interest.

