## Supplemental material for "Coordinated IFN-γ/TNF Axis Drives Selective Loss of Activated Enteric Glia in Inflammatory Bowel Diseases"

### Supplementary information

#### Materials and Methods

<u>Human cohorts used and reanalysed in this study</u>

For the in-house cohort (IBDome), samples were obtained from non-IBD and IBD patients after written informed consent and the ethical committees under the ID EA1/200717 for origin Berlin and 332-17B for origin of Erlangen. Samples from participants undergoing surgery for resection or biopsies were taken during endoscopy at the Gastroenterology Department of the Charité – Universitätsmedizin Berlin, Department of Gastroenterology, Infectious Diseases and Rheumatology as well as the Department of General and Abdominal Surgery | CBF or Universitätsklinikum Erlangen; Medizinische Klinik 1 – Gastroenterologie, Pneumologie und Endokrinologie. Samples were stored in 10 mL RNA protect Tissue Reagent (Qiagen) overnight at 4 °C. For long-term storage, the reagent was removed and biopsies kept at -80 °C. For RNA isolation, the RNeasy Kit (Qiagen) was used according to the manufacturer's instructions employing the TissueLyser LT (Qiagen) and 5 mm steel beads. Concentration and purity of the isolated RNA was determined by the ratios of absorbance (A260:28 and A260:230) using a Nanodrop 1000 (ThermoFisher Scientific). The RNA integrity number (RIN) was determined with the Agilent 2200 Tape Station after preparing the samples as per manufacturer's instructions for the Agilent RNA ScreenTape System. The RNA Clean & Concentrator was used according to manufacturer's instructions for samples with low RIN. Frozen samples were shipped on dry ice to the Quantitive Biology Center (QBIC) of Eberhard Karls Universität Tübingen for sequencing. All patient data has been pseudonymized. The final number of samples analysed from the IBDome cohort were distributed as follows: CD = 277, UC = 121, and non-IBD controls = 46 amounting to a total of 196 samples from male and 198 from female individuals.

For replication studies, analysis was conducted from publicly available bulk and single cell RNASeq datasets from IBD patient cohorts including healthy, Crohn's disease and Ulcerative colitis and the dataset accession numbers or links are provided here below:

Kinchen *et al.* [1], GSE114374 ; Nie *et al.* [2], http://scibd.cn ; Thomas *et al.* [3], GSE282122 ; Oliver *et al.* [4] , https://www.gutcellatlas.org, Mount Sinai Crohn's and Colitis Registry (MSCCR) Argmann *et al.* [5].

Besides these, we also reanalysed previous IBD microarray datasets Ahrens *et al.*, GSE10191 [6]; Gyorffy *et al.* [7] GSE4183; Carey *et al.* [8] GSE9686; Wu *et al*. [9] GSE6731; and Kugathasan *et al.* [10] GSE10616.

<u>Mice and mouse models of intestinal inflammation</u>

All mice on a C57BL/6J background were purchased from Charles River (Charles River GmbH) and Janvier labs (Janvier labs GmbH) and were housed under specific pathogen-free (SPF) conditions. Glia reporter mice were generated by crossing *Plp1*CreERT [11] inducible driver line and the Rosa26-tdTomato mice which carry a loxP-flanked Stop-codon with subsequent fluorescent protein tdTomato cassette in their *Rosa26* locus. Genotyping of the *CreERT* and tdTomato insert was done by polymerase chain reaction on tail DNA. Primer sequences are available upon request. *CreERT*-recombination was induced by oral administration of 75 mg per kg bodyweight of Tamoxifen (Merck KGaA) solubilized stock in ethanol, diluted in corn oil (Merck KGaA).

In similar fashion, *Plp1CreERT* mice were bred with mice carrying a loxP-flanked *Caspase8* allele (*Casp8*$^{flox}$) [12] to generate the *Casp8*$^{i\Delta Glia}$. Genotyping of the *CreERT* and tdTomato insert was done by polymerase chain reaction on tail DNA. Primer sequences are available upon request. The *CreERT*-recombination was induced by oral administration of 75 mg per kg bodyweight Tamoxifen (Merck KGaA) diluted in ethanol and corn oil (Merck KGaA) for three consecutive days.

<u>T-cell activation model with anti-CD3 antibody</u>:

Wildtype C57BL/6J or EGC reporter mice were injected intraperitoneally with 30 µg of purified NA/LE Hamster Anti-Mouse CD3e antibody (BD Bioscience) or Hamster IgG NA/LE Isotype control (BD Bioscience) in PBS, total volume 200 µL. The course of inflammation was monitored by weight, clinical symptoms, and behavioural assessment. Mice were sacrificed after 6 or 24 hours. Disease activity was scored for behavioural changes and clinical symptoms during the aCD3 model. A baseline was set right after the injection for the parameters weight, pilo erection, eyelid ptosis and facial swelling, and activity. Every 3, 6 and 24 hours, the mice were checked and give a score from 0 to 3 for the respective category. A total score for each mouse is reflected by the average of the score value of all categories. The percent weight loss was calculated for a score reflecting 0 to <5% weight loss as score 0, 5% to <10% weight loss as score 1, 10% to <20% weight loss as score 2, and >=20% weight loss as score 3. Pilo erection, eyelid ptosis and facial swelling, and activity were score in increments of 0 to 3. The sample sizes were decided based on previous reports [13].

For the motility assay, the mice were injected with 30 µg of purified NA/LE hamster anti-mouse CD3ε antibody (BD Bioscience) or hamster IgG NA/LE isotype control (BD Biosciences) in PBS, total volume 200 µL, intraperitoneally. After 2 hours, the food was removed from the cage and the mice continued to be monitored. At the five-hour mark, the mice received an oral gavage with 10,000 kDa FITC-Dextran (Merck KGaA) dissolved in 200 µL Lipofundin® MCT20% (B. Braun Melsungen AG). After six hours the mice were sacrificed via cervical dislocation and the digestive tract from stomach to anus removed. After short storage in ice cold PBS, the intestine was laid out on a black background and imaged with the Amersham ImageQuant 800 detector. The intensity of the FITC was measured and quantified with ImageJ and an in-house python script.

The generation and analysis for the following models has been described in detail previously [14]: *Casp8*$^{\Delta IEC}$ ileitis and colitis, *Tnf*$^{\Delta ARE}$ ileitis, acute and chronic dextran sulfate sodium (DSS)-induced colitis, oxazolone-induced colitis, acute and chronic trinitrobenzene sulfonic acid (TNBS)-induced colitis, T-cell transfer colitis, *Eimeria vermiformis* infection, *Helicobacter hepaticus* infection, *Citrobacter rodentium* infection.

Experimental protocols were approved by the Institutional Animal Care and Use Committee of the Regierung von Unterfranken and the Landesamt für Gesundheit und Soziales – Berlin, and in accordance with the UK Scientific Procedures Act of 1986. Generation and analysis of bulk transcriptomic data from the mouse models has described by us recently [14]. The study reference numbers are as follows: 55.2.2-2532-2-1137, -1139, -1178, -1495, -1789; 55.2-2532-2-595, 55.2.2532–2-607, -712; G 0291/18; 54.2532.1.-15/12

<u>Cytokine multiplex</u>

The BioLegend LEGENDplex™ MU Th Cytokine Panel (12-plex) was used to reveal tissue cytokine levels. The tissue was homogenized and the bead assay performed as per manufactures recommendations and instruction. In short, the samples where incubated with the capture beads in V-bottom plate. After washing steps, the detection antibodies were added and in a later step conjugated to the fluorophore. After the final washing steps, the beads were analysed in a BD Accuri C5 flow cytometer. The files were then uploaded to the BioLegend online tool and analysed.

<u>*Ex vivo* plexanoid culture</u>

The isolation of the muscularis externa was adapted from previous protocols [15, 16, 17]. In short, after sacrificing the C57BL/6J, tdTomato reporter, or *Casp8*$^{i\Delta Glia}$ mice, the skin was cleaned with 70% ethanol, the abdominal cavity opened and the small intestine (SI) extracted. Sections of the SI were flushed with ice cold HBSS without calcium and magnesium (Gibco). SI sections of 3 to 4 cm were pulled on sterile knitting needles (3 to 5 mm diameter) and mesenteric fat and endothelial tissue removed. The longitudinal muscle was roughened up with the forceps to create a small incision. With cotton swabs (Cobas Uni Swabs, Roche) the muscularis was peeled off of the SI sections. Muscularis was cut into 5 mm pieces, washed with ice cold HBSS and digested with collagenase type 4 (Worthington) in HBSS with 10% FCS (Merck KGaA) for 1 h at 37°C. Subsequently, single cells were extracted with TrypLE (Gibco) and triturated with 1000 µL pipette tips in culturing medium. Single cells were seeded on Matrigel (diluted 1:30 in sterile PBS) coated plates. For expansion the primary cell culture was done in Neurobasal medium (Gibco) supplemented with 1% FCS (Merck KGaA), 1x B27 (Gibco), 1x Glutamax (Gibco) and Penicilin/Streptamycin (Merck KGaA). After one week, plexanoids were split at a ratio of 1:2 by releasing the single cells with TrypLE (Gibco) and seed them on new Matrigel (Corning) coated cell culture containers for further experiments. Further expansion was done again in supplemented Neurobasal medium.

<u>Stimulation of *Casp8*$^{i\Delta Glia}$ plexanoids</u>

As described before, plexanoids of the *Casp8*$^{i\Delta Glia}$ mouse line were isolated and expanded, then replated on Matrigel (Corning) coated (diluted 1:30 in sterile PBS) 48-well cell culture plates. After 4 days of further proliferation in this setup, the medium was changed to Neurobasal medium (Gibco) supplemented with Glutamax (Gibco) and Penicillin/Streptomycin (Merck KGaA) including a vehicle control versus 50ng/mL recombinant mouse TNF. For live cell imaging, 5µg/mL propidium iodide was added to the medium and the vessels monitored with the Incucyte live cell imager (Sartorius GmbH) for 72 hours.

<u>Sorting, culturing and stimulation of enteric glial cells</u>

Muscularis externa was isolated as described before from Glia reporter mice. In the first culturing period, cells were cultured like plexanoids to expand the population of tdTomato positive glia cells. Subsequently, the expression of tdTomato was induced by adding 500nM (Z)-4-hydroxytamoxifen (Merck KGaA) directly to the cell culture medium for three consecutive days. Finally, single cells were obtained with TrypLE (Gibco) and transferred in FACS buffer consisting of PBS with 1 % FCS (Merck KGaA) and 2 mM EDTA (Gibco). The sorting of tdTomato positive Glia cells was done by the FACS Core Unit of the Friedrich-Alexander-Universität Erlangen-Nürnberg (FAU). After spinning down the sorted EGC population, cells were seeded on plates coated with Matrigel (Corning) diluted 1:30 in sterile PBS and grown in Neurobasal medium (Gibco) supplemented with 1% FCS (Merck KGaA), 1x B27 (Gibco), 1x Glutamax (Gibco), Penicilin/Streptomycin (Merck KGaA), 20 ng/mL GDNF (Peprotech/ThermoFisher Scientific) and 20 ng/mL FGF-basic

(Peprotech/ThermoFisher Scientific). For cytokine stimulations, cell culture medium was substituted to Neurobasal medium (Gibco) supplemented with Glutamax (Gibco), Penicilin/Streptomycin (Merck KGaA), 20 ng/mL GDNF (Peprotech/ThermoFisher Scientific) and 20 ng/mL FGF-basic (Peprotech/ThermoFisher Scientific) with the respective cytokine at 50 ng/mL (See supplemental table ST1).

**Supplemental table ST1: Cytokine and other stimulant concentrations for EGC cell culture**

| Cytokine | Concentration | Catalogue # | Company |
|---|---|---|---|
| Ms TNF recombinant | 50 ng/mL | 12343016 | ImmunoTools GmbH |
| Ms IFN-γ recombinant | 50 ng/mL | 575306 | BioLegend, Inc. |
| Ms IL-1β recombinant | 50 ng/mL | 12340013 | ImmunoTools GmbH |
| Ms Il-17A recombinant | 50 ng/mL | 12340174 | ImmunoTools GmbH |
| zVAD fmk | 20 µM | S7023 | Selleckchem |
| Ms GDNF recombinant | 20 ng/mL | 450-44 | PeproTech/ ThermoFisher Scientific |
| Hu FGF-basic recombinant | 20 ng/mL | 100-18B | PeproTech/ ThermoFisher Scientific |

<u>Cell death in stimulated, sorted enteric glia cells</u>

For the induction of cell death, sorted EGC were stimulated for 24 hours in reduced medium with IFN-γ (BioLegend, Inc.), TNF (ImmunoTools GmbH), Z-VAD-FMK (Selleckchem) and combinations of these drugs (see supplemental table ST1). For the live cell imaging CytoxGreen (Sartorius GmbH) was added according to the manufacturer's recommendation. At the end of the experiment, the cells were used for immunostaining using anti-mouse phosphorylated Ser345 MLKL. For the immunostaining of the stimulated glia, the medium was exchanged to 4% PFA and incubated for 10 minutes at room temperature (RT). After washing twice with PBS, the cells were permeabilized with 1% TritonX (Merck KGaA) in PBS for 10 minutes at RT. Subsequent blocking was performed for 30min at RT in PBS containing 1% BSA (Carl Roth GmbH + Co. KG), 10% FCS (Merck KGaA) and 0,025% TritonX (Merck KGaA). The same buffer was used for the primary antibody (overnight at 4 °C) and secondary antibody (2 hours at RT) incubation with two PBS wash cycle after each incubation. Finally, Hoechst was added for nucleus counter staining for 10 minutes at RT. The images were acquired with a Leica SP5 confocal microscope.

**Supplemental table ST2: Antibody usage and source information**

| Antibody | Type | Dilution | Species | Code | Company |
|---|---|---|---|---|---|
| Primary antibody | | | | | |
| Anti-S100B | Unconjugated | 1:2 | Rabbit | GA504 | Agilent Dako |
| Anti-TUBB3 | Biotinylated | 1:100 | Mouse | 801212 | BioLegend, Inc. |
| Anti-GFAP | Alexa Flour 488 | 1:400 | Mouse | 644704 | BioLegend, Inc. |
| Anti-CD4 | Unconjugated | 1:100 | Rat | BD (553043) | BD Biosciences |
| Anti-pMLKL (S345) | Unconjugated | 1:400 | Rabbit | 37333S | Cell Signaling Technologies |
| Secondary antibody | | | | | |
| Anti-Rabbit | Biotinylated | 1:500 | Goat | 111-065-144 | Jackson |
| Anti-Rat | Biotinylated | 1:500 | Goat | 554014 | BD Biosciences |
| Anti-Rabbit | Alexa Flour 647 | 1:400 | Donkey | 406414 | BioLegend, Inc. |
| Anti-Rat | Dylight 550 | 1:400 | Donkey | SA5-10027 | ThermoFisher Scientific GmbH |
| FACS antibodies | | | | | |
| Anti-GFAP | Brilliant Violet 421 | 1:200 | Mouse | 644710 | BioLegend, Inc. |

| | | | | | |
|---|---|---|---|---|---|
| MHC II I-A/E-A | FITC & APC-Cy7 | 1:2000 | Rat | 107606 & 107628 | BioLegend, Inc. |
| Western blot antibodies | | | | | |
| PLP1 | Unconjugated | 1:500 | Mouse | MCA839G | BIO-RAD |
| ACTB | HRP conjugated | 1:2500 | Mouse | ab49900 | Abcam |

<u>Flow cytometry analysis of enteric glia population dependent cell death.</u>

In case of *in vivo* cell death FACS analysis, muscularis externa was stripped from intestines of tdTomato positive mice after 6h of aCD3 or isotype antibody injection as described before. The muscularis externa was then cut in small pieces and digested in DMEM/F12 (Gibco) with Liberase TH (Roche), Dispase (Merck KGaA) and DNase (Merck KGaA) for 30 minutes at 37°C. After following trituration in DMEM/F12 (Gibco) with 2.5% BSA (Carl Roth GmbH + Co. KG) and 5 mM EDTA (Gibco), the cells were resuspended in FACS buffer (PBS with 1% FCS, 2mM EDTA) and filtered through 70 and 30 µm cell filters (Corning).

Subsequently the respective cell solution was centrifuged at 400 rcf for 7 minutes at 4°C. The supernatant was discarded and the cell pellet resuspended in FACS buffer. The different conditions were then stained with a fixable Live/Dead stain (ThermoFisher Scientific GmbH) and MHC class II antibody (BioLegend, Inc.) for 45 minutes at 4°C. After washing, cells were fixed in 4% PFA for 15 minutes at room temperature, washed again in FACS buffer, then stained for GFAP (BioLegend, Inc.). Flow cytometry was performed with the BD Fortessa system, the data analysed in FlowJo v10.10.

<u>H&E-stained paraffin sections of intestinal tissue were used for muscle thickness measurements.</u>

The muscle thickness was defined as the perpendicular distance from the bottom of the crypts to the outer gut wall and was measured using the NDP.view2 software. Areas containing lymphoid follicles and Payer’s patches, or where the section ran longitudinal to the axis of the gut were excluded from the analysis. All clearly defined crypts were chosen and the final quantification is based on average distances from one section with an average of ~40 crypt-muscle measurements per mouse. Statistics are drawn on biological replicates, not on number of crypts.

Immunohistochemistry and Immunofluorescence staining

For the haematoxylin and eosin (H&E) staining, we prepared sections of 5-10 μm from tissues fixed in formalin and embedded in paraffin, which were then subjected to standard H&E staining protocol. H&E images were obtained using a NanoZoomer 2.0 (Hamamatsu).

The H&E scoring for bleeding is defined by the amount of red blood cells clearly outside of blood vessels and in the lumen of the intestine. The H&E sections were taken from the same duodenal region for each animal, and the isotype control animals used for baseline comparison. The scoring of *Casp8*$^{\text{flox}}$ animal and *Casp8*$^{\text{i}\Delta\text{Glia}}$ treated with anti-CD3 was performed blind with anonymized filenames.

The immunostainings were performed on paraffin sections of the tissues, which were cut, deparaffinized, hydrated, and treated with a Tris-EDTA-based antigen retrieval solution. To prevent non-specific binding, a commercial blocking reagent (Immunoblock 1X, Carl Roth GmbH + Co. KG) was utilized. The tissue sections were then incubated overnight at 4°C in the dark with the primary antibody for each immunostaining (Supplemental table ST2). We washed the samples in TBS and then we incubated them with either the secondary antibody or streptavidin conjugates (DyLight, ThermoFisher Scientific GmbH) for 1h at 4°C. Samples then were counterstained with Hoechst 33342 (ThermoFisher Scientific GmbH) for the cellular nuclei and cover-slipped in fluorescence mounting medium (Agilent Dako). A Leica TCS SP5 confocal microscope (Leica Microsystems) was employed for the acquisition of the immunofluorescence images using the necessary settings.

Whole mount immunostaining

Whole mount staining was performed as previously described, with some modifications. Intestinal tissue was obtained from duodenum and jejunum in approximately 1 cm length, opened longitudinally and placed between two microscopy slides. These slides were submerged in 4% ice cold PFA and carefully agitated for no longer than 2 hours. After washing the tissue twice with PBS, 0.025% Triton X (Merck KGaA) and 0.02% Tween20 (Merck KGaA) in PBS was used to permeabilize the tissue. After blocking unspecific binding side with Roti Histoblock (Carl Roth GmbH + Co. KG) in 0.025% Triton X (Merck KGaA) and 0.02% Tween20 (Merck KGaA) in PBS, primary antibodies were incubated for 7 days at 4°C in a blocking solution with Roti Histoblock (Carl Roth GmbH + Co. KG), 0.0125% Triton X (Merck KGaA), 0.01% Tween20 (Merck KGaA) and 0.01% sodium azide (Carl Roth GmbH + Co. KG) in PBS. After subsequent washing steps with PBS, secondary antibodies were incubated in blocking solution for 4 days. After final washing steps in PBS, tissue was stored in PBS containing 0.02% sodium azide (Carl Roth GmbH + Co. KG). For imaging, the tissue was placed between two coverslips with mounting medium. Imaging was performed on a Leica confocal microscope SP5.

<u>RNA isolation, reverse transcription, and qPCR</u>

RNA was obtained from homogenized tissue or FACS sorted EGC with the column-based RNeasy kit (Macherey-Nagel GmbH). After quality control via Nanodrop, 200 ng of the RNA was processed into first strand cDNA using the reverse transcriptase kit (Jena Bioscience GmbH) according to manufactures instructions. For qPCR, 2 µL of cDNA was used in a 20 µL SYBR Green based PCR reactions in 96-well format, reaction performed in a BioRad PCR CFX Connect OM.

<u>Bulk mRNA Sequencing</u>

RNA was obtained as described before. RNA quality control was performed with Nanodrop analysis and Agilents TapeStation automated electrophoresis. Between 100 to 200 µg of total RNA samples were shipped to and analysed by Novogene GmbH, Munich, Germany, including a secondary quality control and library preparation. Sequencing results were analyzed with an in-house pipeline described earlier [14].

<u>Western blotting</u>

Protein samples were prepared from snap-frozen tissue. The samples were put in ice-cold T-Per lysis buffer (Merck KGaA) with protease and phosphatase inhibitors (Roche, 1 tablet per 10mL) and 1% PMSF (Roche). With a steel bead added to the tissue, the sample was homogenized in a high-speed shaker (Retsch GmbH) at 25 Hz for 2 minutes. Further lysis was performed on ice for 20 minutes and subsequent centrifugation for 20 minutes at 4°C, 14.000 rpm. Protein concentrations from the supernatants were determined via standard Bradford method and equal amounts of protein per sample were mixed with 4x loading buffer containing 40 mM 1,4-Dithiothreit (DTT, Carl Roth GmbH + Co. KG). The samples were boiled at 95°C for 5 minutes and loaded on a pre-cast SDS-PAGE-gels (BIO-RAD).

Gels were run in BioRad electrophoresis chambers at 200V for approximately 45 minutes. The gel was transferred on PVDF () membranes with the BioRad Turboblotter and membranes blocked with 5% milk powder (Merck KGaA) for 1h at room temperature. Subsequently the membranes were probed for the target proteins with primary antibodies (Supplemental table ST2) diluted in 5% BSA (Carl Roth GmbH + Co. KG) in TBST overnight at 4°C. Secondary antibodies coupled with HRP are added after washing and incubated for 2 hours at room temperature. Signal detection with Western Lightning Plus ECL (Revvity, Inc.) was done with the Amersham ImageQuant 800 detector. Total change of proteins was normalized to β-Actin. Membranes that were probed for more than one protein were stripped-off the bound antibodies with a gentle stripping buffer consisting of 1 g/L SDS (Carl Roth GmbH + Co. KG), 15 g/L Glycine (Carl Roth GmbH + Co. KG), 10 mL of Tween 20 (Merck KGaA), brought to a pH of 2.2 with 32% HCl (Carl Roth GmbH + Co. KG). The membranes were shaken in the gentle stripping buffer twice for 10 minutes at RT, washed with PBS and TBST for a total four times for 5 minutes, two times respectively.

Single nucleus RNA sequencing of formalin-fixed paraffin embedded tissue samples

Human formalin-fixed and paraffin-embedded (FFPE) full-thickness intestinal wall tissue sections of resected tissue were processed as per the kits manufacturer instructions (‘CG000632_RevC’ 10x Genomics, Single Cell Gene Expression Flex Kit). In short, the tissue was deparaffised and rehydrated. After transfer to ice-cold PBS the tissue was dissociated using a pellet pestle in a dissociation enzyme mix. After trituration and filtering through a 30µm filter, the single nuclei were collected and counted. After quality control of the nuclei, probe hybridization was performed for the nuclei in bulk. Subsequently the nuclei were packed into GEMs with the 10x Genomics droplet-based single cell cassette. Inside the GEMs, probes were first ligated and subsequently received a specific barcode to identify the single cells after sequencing. Probe hybridization and library construction were carried out as described in User Guides ‘CG000527_Rev_E’ for multiplexed samples or ‘CG000691_Rev_A’ for singleplex samples targeting 10.000 cells per sample. Sequencing was performed in the genomics core facility of the Max Delbrück Center - Berlin Institute for Medical Systems Biology (MDC-BIMSB) with the Illumina Nova Seq X targeting 20.000 read pairs per cell.

Demultiplexing of sequenced libraries was achieved with bcl2fastq (Illumina). FASTQ data processing was done with the multi pipeline from Cell Ranger (version 7.1.0, 10x Genomics), utilizing the reference genome GRCh38-2020-A for alignment. Analyses were performed in R (version 4.3.3) with the Seurat package (version 5.0.1). At first, ambient RNA was removed using SoupX (version 1.6.2) and subsequently, all samples were filtered to comprise only cells with more than 400 genes per cell, more than 450 UMIs per cell, a fraction of mitochondrial reads lower than 15% and a fraction of haemoglobin reads lower than 5%. Thereafter, doublets were excluded with Doublet Finder (version 2.0.4) and the gene expression data for all samples was merged. An additional filtering step was carried out to remove cells with more than 4.000 genes and more than 10.000 UMIs per cell. Next, log normalization, scaling, PCA and UMAP were performed selecting the first 35 PCs for downstream analysis. Harmony (version 1.2.0) served to integrate the dataset by sample followed by SNN-graph based clustering (resolution = 0.25). Cell types were first annotated based on marker genes from literature into epithelial cells (EPCAM, KRT8), stromal cells (CDH5, COL1A1, COL1A2, COL6A2, VWF) or immune cells (PTPRC, CD3E, CD3G, CD3D, CD79A, CD79B). Thereafter, each compartment (epithelial, stromal, immune) was subclustered respectively and manually annotated to obtain more detailed levels of cell type information. Finally, all annotation levels were mapped back onto the original object. This dataset is part of an ongoing study, therefore for this study, the enteric glia cells of the dataset were extracted as a standalone file with the typical pre-processing was done seperately.

Combined cell death signature gene list

To track the changes in cell death signature genes, we used a combined approach tracking human and mouse genes belonging to the KEGG pathways for both apoptosis and necroptosis. Several of these genes share regulatory behaviour and tend to be coregulated with each other. Therefore, we generated the combined cell death signature list from the pathway gene lists shown in supplemental table ST3 below. The average expression of these genes was tracked on the respective human and mouse snRNA-Seq and scRNA-Seq datasets.

**Supplemental table ST3: Combined cell death pathway genes for mouse and human**

| **Mouse** | **Human** |
|---|---|
| Aifm1 | AIFM1 |
| Anxa1 | ANXA1 |
| Apaf1 | APAF1 |
| Bad | BAD |
| Bax | BAX |
| Bcl2 | BCL2 |
| Bgn | BGN |
| Birc2 | BIRC2 |
| Birc3 | BIRC3 |
| Casp1 | CASP1 |
| Casp3 | CASP10 |
| Casp10 | CASP3 |
| Casp4 | CASP4 |
| Casp6 | CASP6 |
| Casp7 | CASP7 |
| Casp8 | CASP8 |
| Cyld | CYLD |

| | |
|---|---|
| Egr3 | EGR3 |
| Emp1 | EMP1 |
| Etf1 | ETF1 |
| F2r | F2R |
| Fas | FAS |
| Fasl | FASLG |
| Gch1 | GCH1 |
| Hgf | HGF |
| Ier3 | IER3 |
| Ifnar1 | IFNAR1 |
| Ifnar2 | IFNAR2 |
| Ifngr1 | IFNAR3 |
| Il1b | IFNGR1 |
| Il1rap | IL1B |
| Il6 | IL1RAP |
| Irf1 | IL6 |
| Isg20 | IRF1 |
| Lef1 | ISG20 |
| Mlkl | LEF1 |
| Nfkb1 | MLKL |
| Nfkbia | NFKB1 |
| Ntrk1 | NFKBIA |
| Pdgfrb | NTRK1 |
| Pmaip1 | PDGFRB |
| Psen2 | PMAIP1 |
| Rela | PSEN2 |
| Ripk1 | RELA |
| Ripk3 | RIPK1 |
| Smpd1 | RIPK3 |
| Sod2 | SMPD1 |
| Tap1 | SOD2 |
| Tnfrsf10b | TAP1 |
| Tnfrsf1a | TNFRSF10A |
| Tnfrsf12a | TNFRSF12A |
| Tradd | TNFRSF1A |
| Traf2 | TNFRSF10B |
| Wee1 | TRADD |
| Xiap | TRAF2 |
| Zbp1 | WEE1 |
| | XIAP |
| | ZBP1 |

Plot generation software

Data was analysed in commercial software GraphPad Prism program (Version 10) or in-house python (Version 3.11.0) and R scripts (Version 4.4.0). For heatmaps the ComplexHeatmap (Version 2.20.0) package was used, Ridge plots were generated using the ggridges package (0.5.6). Other plots in R were generated using the ggplot2 package (3.5.1). Matplotlib (3.9.0) and Seaborn (0.13.2) were used for the figure plots generated using python (3.11.0). Other packages used for data analysis and visualization are listed in supplemental table ST4 below.

**Supplemental table ST4: Software packages and versions used in data analysis and visualization**

| Package name | Version |
|---|---|
| **R version 4.3.3** | |
| ggplotify | 0.1.2 |
| progeny | 1.24.0 |
| ggfortify | 0.4.16 |
| RColorBrewer | 1.1-3 |
| ComplexHeatmap | 2.18.0 |
| ggrepel | 0.9.5 |
| scales | 1.3.0 |
| openxlsx | 4.2.5.2 |
| tidyr | 1.3.0 |
| ggplot2 | 3.4.4 |
| cowplot | 1.1.2 |
| DESeq2 | 1.42.0 |
| extrafont | 0.19 |
| fmsb | 0.7.6 |
| dplyr | 1.1.4 |
| tidyverse | 2.0.0 |
| ggridges | 0.5.5 |
| decoupleR | 2.8.0 |
| gridExtra | 2.3 |
| circlize | 0.4.16 |
| **Python 3.12.3** | |
| numpy | 1.26.2 |
| pandas | 2.0.3 |
| scanpy | 1.10.4 |
| anndata | 0.11.1 |

| | |
|---|---|
| matplotlib | 3.8.2 |
| scipy | 1.11.4 |
| seaborn | 0.13.2 |
| scipy | 1.11.4 |
| statsmodels | 0.14.4 |
| igraph | 0.11.8 |

#### Supplementary figure legends

**Supplemental Figure 1. Correlation of canonical and novel enteric glial cell (EGC) markers on in-house and replication cohort.** **A & B)** Correlation plots for the patient-wise transcript expression levels (each dot represents levels from one patient) of the indicated inflammation markers (*TNF*, *SAA1*) versus canonical EGC markers *S100B*, and *PLP1* across all samples in the IBDome cohort irrespective of sampling procedure or degree of inflammation for UC (A) and CD (B). **C)** Expression levels of EGC enriched genes obtained as per scheme in Fig. 1A, on six indicated publicly available microarray datasets from IBD patients showing consistent reduction of *PLP1* (arrows) **D)** Representative photomicrographs of immunostained tissues showing high levels of mucosal S100B positive cells from active UC patients compared with non-IBD controls. **E)** Network plot where nodes are genes (beige) or histology scores (red, riley = modified median Riley score for UC and naini = modified median Naini Cortana score for CD) from colonic biopsy samples of the IBDome cohort. Edge lengths between nodes represent correlation with edge color indicating direction of correlation (green = positive, red = negative), and edge thickness indicating correlation strength. **F & G)** Heatmaps comparing mean z-scores calculated from normalized counts for indicated EGC enriched genes across IBD patients from the IBDome (F) and the Mount Sinai Crohn's and Colitis Registry (MSCCR) cohorts (G) from ileal (Ile_) and colonic (Col_) biopsies or resected samples with red dots indicating genes of interest. Asterisks indicate p-values $< 0.05$ from Mann-Whitney U test applied on normalized counts.

**Supplementary Figure 2. Transcriptomes from IBD single enteric glial cells (EGC) and single EGC nuclei reveal cluster and IBD subtype-specific induction in cell death and activation signatures.** **A)** Uniform manifold approximation and projection (umap) plots from single nuclei mRNA sequencing from colonic full thickness formalin-fixed and paraffin embedded tissue sections from IBD patients and controls. The EGC cluster is circled with dashed line **B)** Dot plot showing the identification of the EGC cluster using glia enriched markers. **C)** Violin plots showing disease type-wise quantification of indicated genes or pathway signatures reanalysed from the dataset reported by Oliver *et al*.[4]. Abbreviations for disease subtype are: CD = Crohn's disease, Neig-Inf = neighbouring inflamed, Pedi-IBD = paediatric IBD, UC = ulcerative colitis. The numbers on the plots indicate FDR corrected p-values derived from the two-tailed Mann-Whitney U test versus healthy controls, ns = not significant. D **& E)** Thresholded expression levels for the indicated genes and for cell death signature together with a blend showing UC specific elevation in interferon target *CXCL9,* and of the cell death pathway signature from EGCs on the Kinchen *et al*. [1] scRNA-Seq dataset (D) and from the integrated IBD scRNA Seq dataset by Nie *et al.* [2] (E). **F & G)** umap plots showing the expression levels of indicated genes and that of the cell death signature along with the inflammation status per cell (Inf_status) from UC (F) and CD (G) from the single EGC transcriptomes reanalysed from the dataset by Thomas *et al.* [3]. **H)** Violin plots showing quantitative difference for indicated genes and cell death pathway signature reanalysed from the Thomas *et al.* [3] dataset shown in (F) and (G) above. The numbers on the violin plots indicate FDR corrected p-values derived from the two-tailed Mann-Whitney U test versus healthy controls.

**<u>Supplemental Figure 3. Expression of canonical enteric glial cell (EGC) markers in colitis mouse models and reduced PLP1 protein in T-cell-driven gut inflammation model</u> A)** Radar charts showing log2 fold changes of key mouse EGC genes in different paradigms of modelling colitis [DSS = dextran sulfate sodium induced colitis; TNBS = trinitrobenzene sulfonic acid-induced colitis; Oxa = Oxazolone-induced colitis; T-transfer = adoptive T-cell transfer-induced colitis; Casp8(col) = epithelial specific ablation of *Casp8* -induced colitis, C.rode = *Citrobacter rodentium* infection-induced colitis; H.hepa = *Helicobacter hepaticus* infection and IL-10R inhibition-induced colitis; Wound = endoscopic pinch-biopsy wound-induced colon injury, h6 and h48 indicate 6 and 48 hours post-biopsy injury] [14, 18]. Asterisks indicate p-values < 0.05 from the DESeq2 Wald statistics test, grey hexagon outlines scale for log2 fold change at zero **B**) Volcano plot showing extraction of EGC enriched markers from previously published sorted EGC vs non-EGC cells [19] **C & D)** Detection and quantification of muscularis inflammation in the anti-CD3 antibody antibody induced intestinal inflammation model (aCD3) (C) Representative H&E image showing quantification method of muscle wall thickness from crypt base to serosa and (D) comparison of muscle wall thickness in micrometres from the tissues of aCD3 versus isotype (Iso) treated mice at the indicated time points from three to four independent biological replicates per group,

Asterisks indicate $p < 0.05$ from non-parametric, Mann-Whitney test. **E)** Levels of indicated cytokines measured from duodenum of mice treated with isotype and aCD3 groups from the indicated time-points. Data are derived from three to four biological replicates per group. Asterisks indicate $p < 0.05$ from one-way ANOVA followed by the Kruskal-Wallis post-hoc test **F)** Quantification of PLP1 protein levels of whole tissue in the indicated groups via western blotting. Asterisks indicates $p < 0.05$ in a two tailed unpaired Student's t-test $n = 3$.

**Supplementary Figure 4. Sorting of enteric glial cells (EGC) and confirmation of type-1 immune-related activation in EGC clusters from previous datasets. A)** Gating strategy for fluorescence-activated cell sorting of tdTomato+ EGC from the muscularis of glia reporter mice. **B)** Thresholded expression densities of several activation-associated genes from the indicated clusters of mouse single EGC transcriptomes reported by Progatzky *et al.* [19].

**Supplementary Figure 5: Induction of cell death in activated enteric glial cells (EGC) *in vivo*. A)** Plots showing gating strategy for tdTomato+ single EGC from digested muscularis samples for further analysis of **B)** activated (tdTomato+, GFAP+) live and dead (fixable dead cell dye+) EGC and their proportions in aCD3 and isotype (Iso) groups.

**Supplementary Figure 6: Coordinated action of IFN-γ and TNF drive EGC activation, necroptosis, and repress enteric gliogenesis A)** Schematic depiction of the protocol for the establishment of FACS sorting tdTomato+ EGC and culturing them *ex vivo* for transcriptomic and live cell imaging evaluation. **B)** Representative photomicrographs of sorted EGC showing tdTomato reporter expression, traceable in live culture **C-E)** Representative photomicrographs from sorted EGC, fixed and immunostained for the canonical EGC marker SOX10 (C) and S100B (D) showing that all sorted tdTomato+ cells are positive for SOX10 and S100B with some expressing the neuronal maker TUBB3 (E) after 10 days in *ex vivo* culture **F & G)** Volcano plot showing transcriptomic response of sorted EGCs to IL-1β (F) and IL-17A (G) stimulation *ex vivo*. Data is derived from three independent biological replicates per group, dashed brown line along the x-axis denotes adjusted p-value threshold of 0.05 for the DESeq2 Wald statistics test. Vertical dashed brown lines indicate absolute log2 fold change threshold of 0.58. **H & I)** Scatter plots showing transcriptome-transcriptome comparison of shared genes between TNF versus IFN-γ stimulated (H) and IL-1β versus TNF stimulated (I) sorted tdTomato+ EGC. **J & K)** Selected upregulated (J) and downregulated (K) ontologies from the transcriptomes of sorted EGC co-stimulated with IFN-γ and TNF from Fig.6C. **L)** Representative photomicrographs of immunostained plexanoids cultured from intestines of C57BL/6J mice displaying S100B positive

EGC surrounding TUBB3 positive neurons, displaying morphologies reminiscent of plexuses in culture. **M)** Heatmap visualizing normalized counts of selected cell type marker genes from bulk RNASeq delineating the composition of plexanoid cultures from three independent biological replicates (M = mouse, M1 to M3) showing dominance of EGC in the culture. **N - P)** Mean with standard deviation of disease activity scores (N), muscle thickness (O) and H&E bleeding scores (P) from littermate mice with glia sufficient in *Casp8* (*Casp8*$^{flox}$) and glia with an inducible deficiency of *Casp8* (*Casp8*$^{i\Delta Glia}$) mice treated with 75mg per kg body weight of tamoxifen for 5 days followed by the induction of intestinal inflammation via the aCD3 model. Data is derived from four independent biological replicates. Asterisks in (P) indicate $p < 0.05$ in Welch's two tailed t-test.
