## Supplementary figures and images for "Coordinated IFN-γ/TNF Axis Drives Selective Loss of Activated Enteric Glia in Inflammatory Bowel Diseases"

### Supplemental Figure 1

#### Supplemental Figure 1

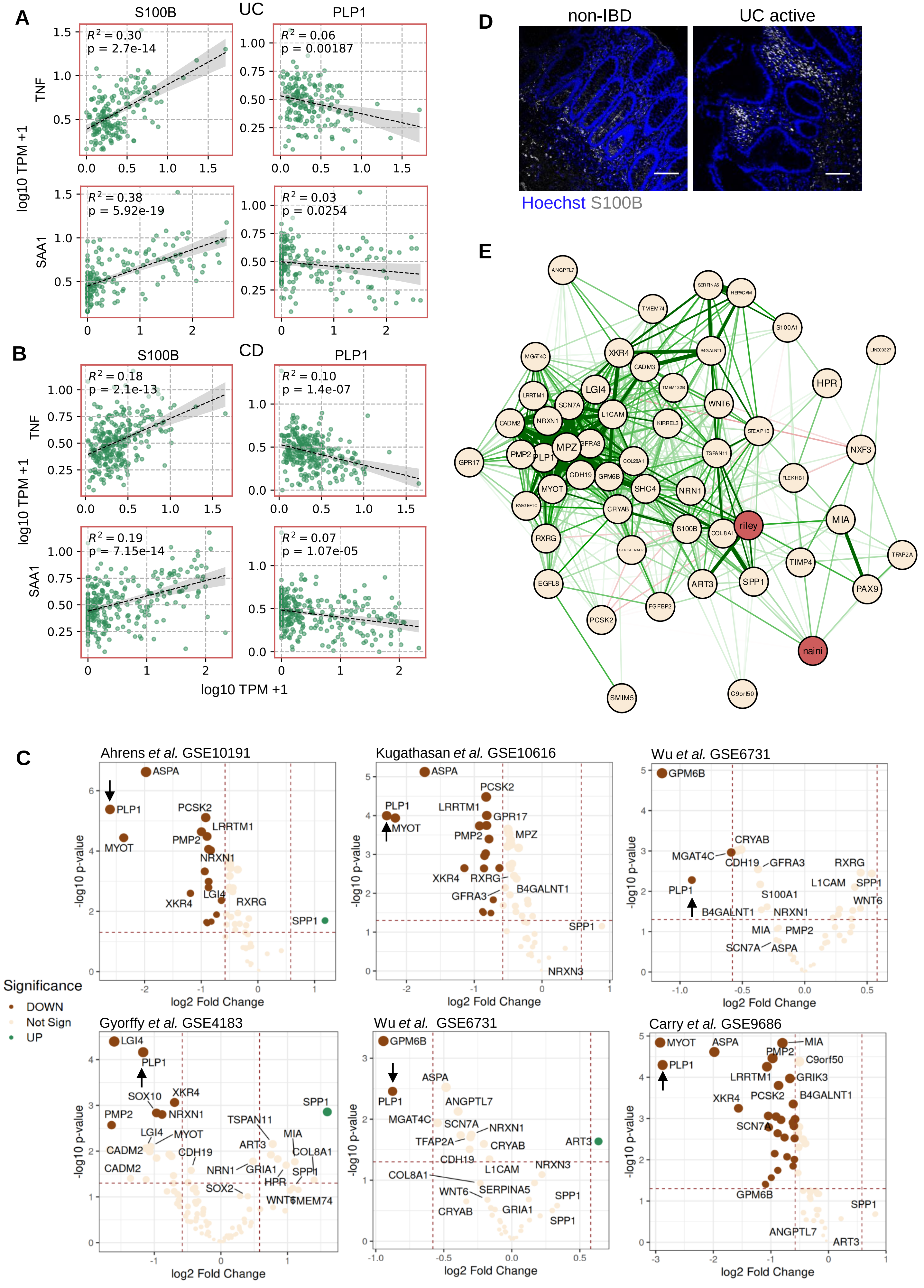

#### Supplemental Figure 1

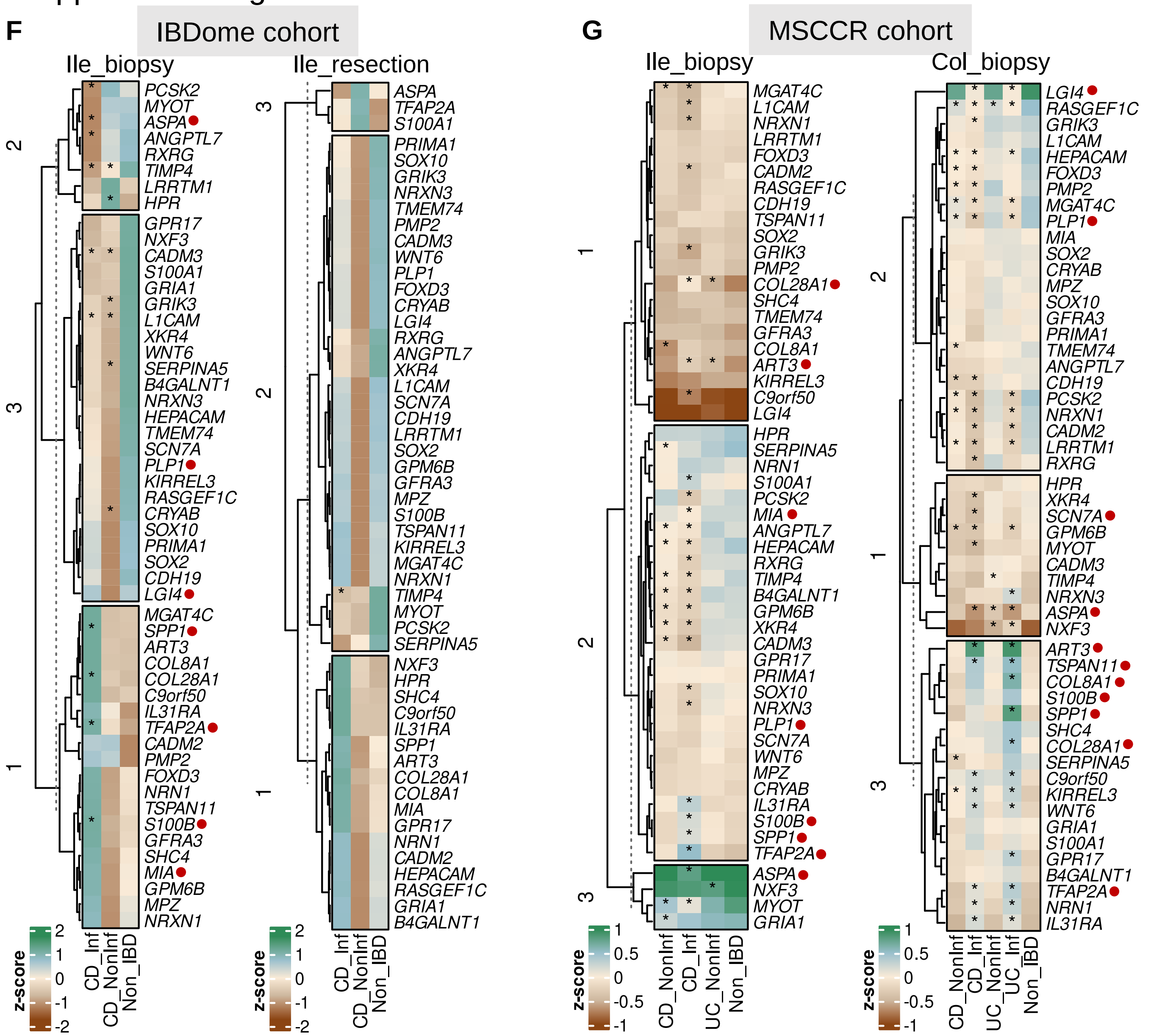

### Supplemental Figure 2

#### Supplemental Figure 2

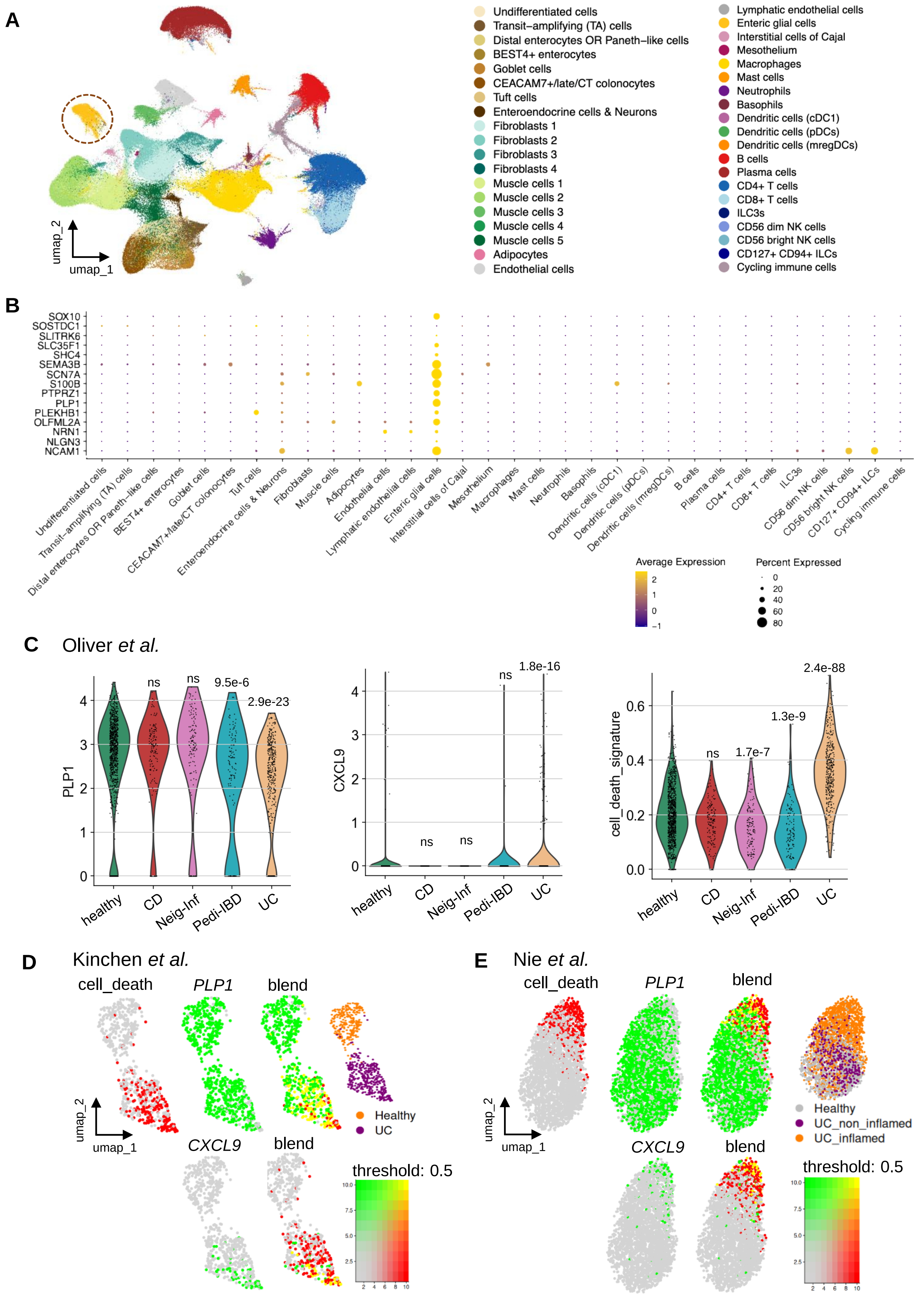

Supplemental Figure 2

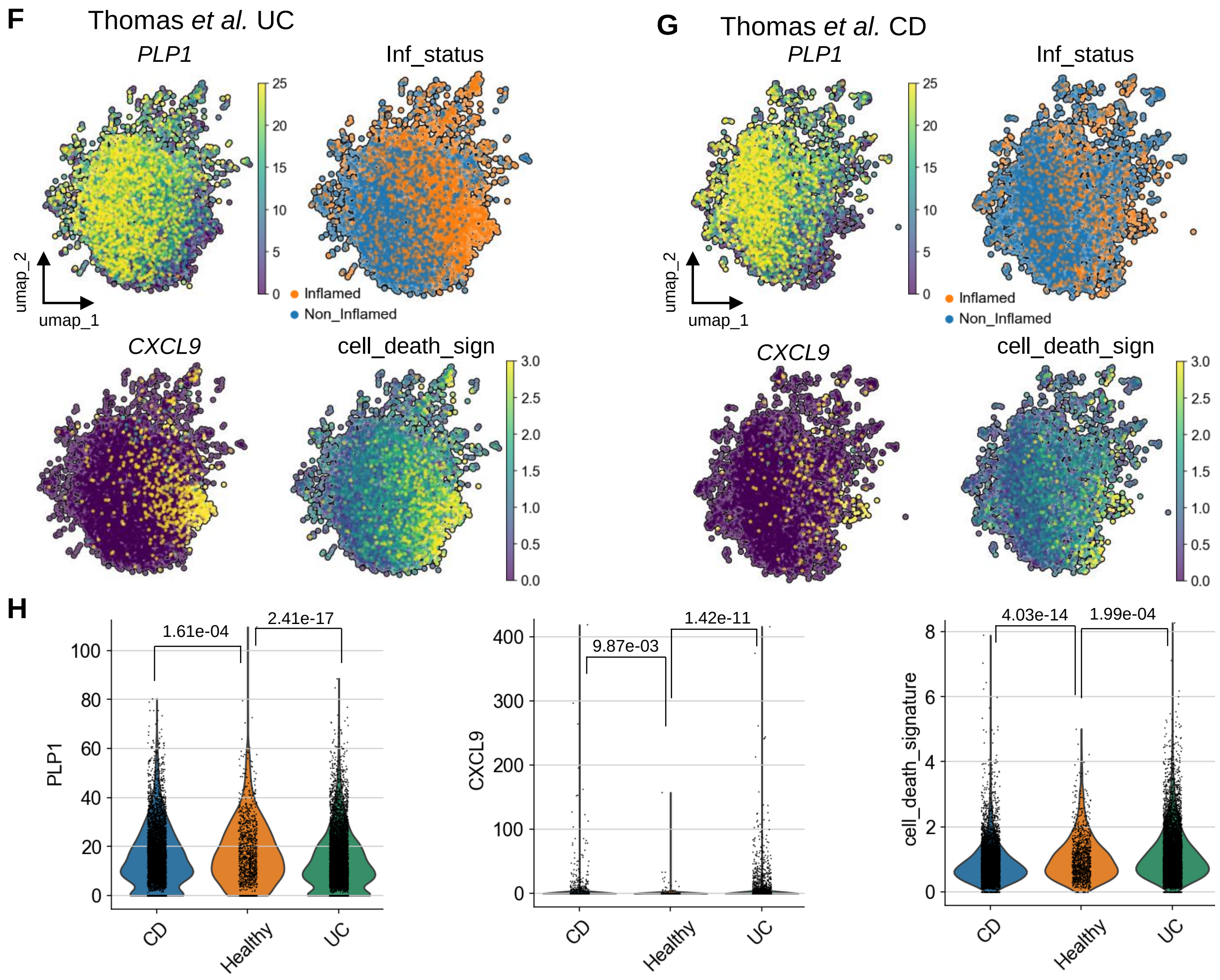

### Supplemental Figure 3

#### Supplemental Figure 3

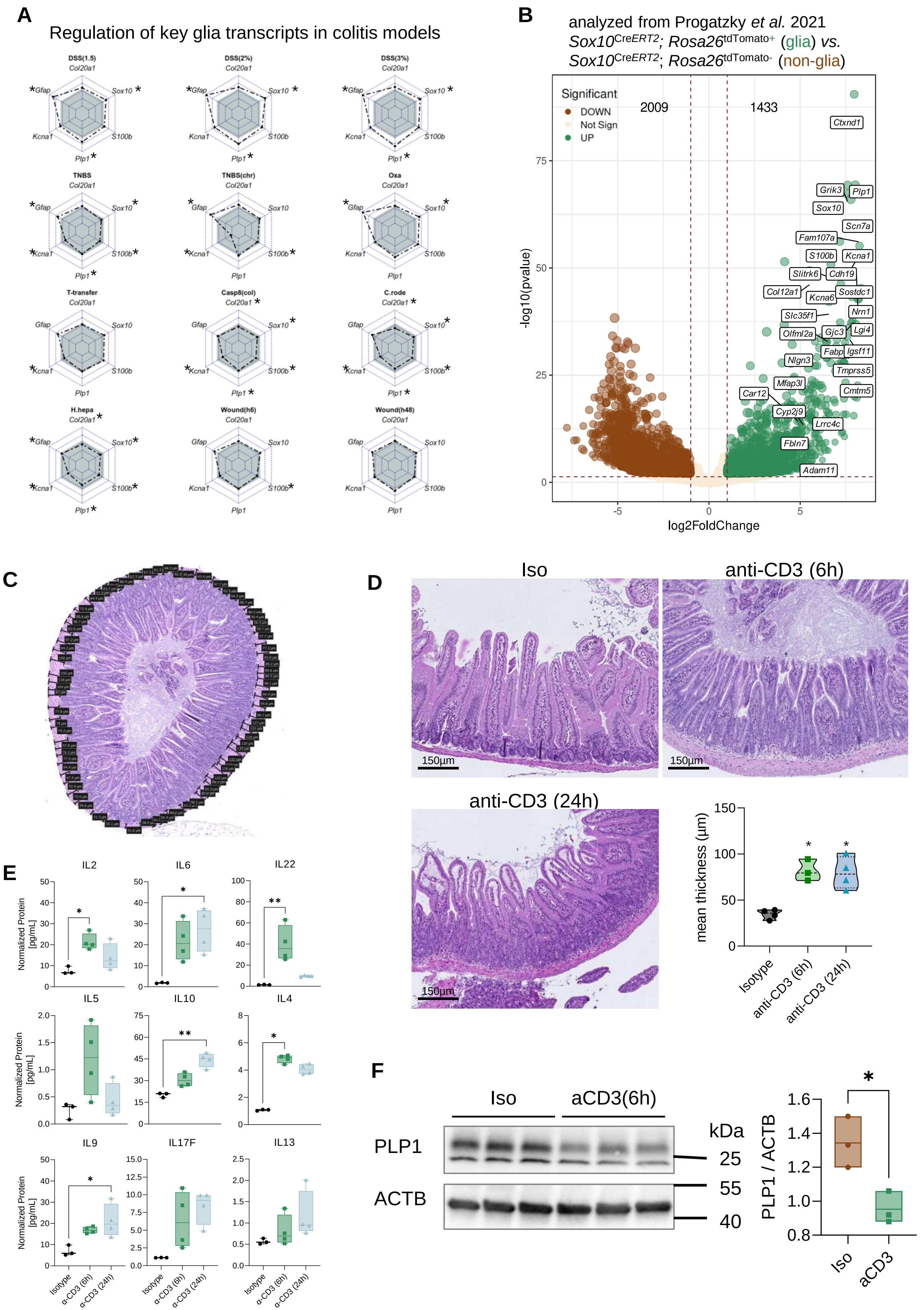

### Supplemental Figure 4

#### Supplemental Figure 4

**A**

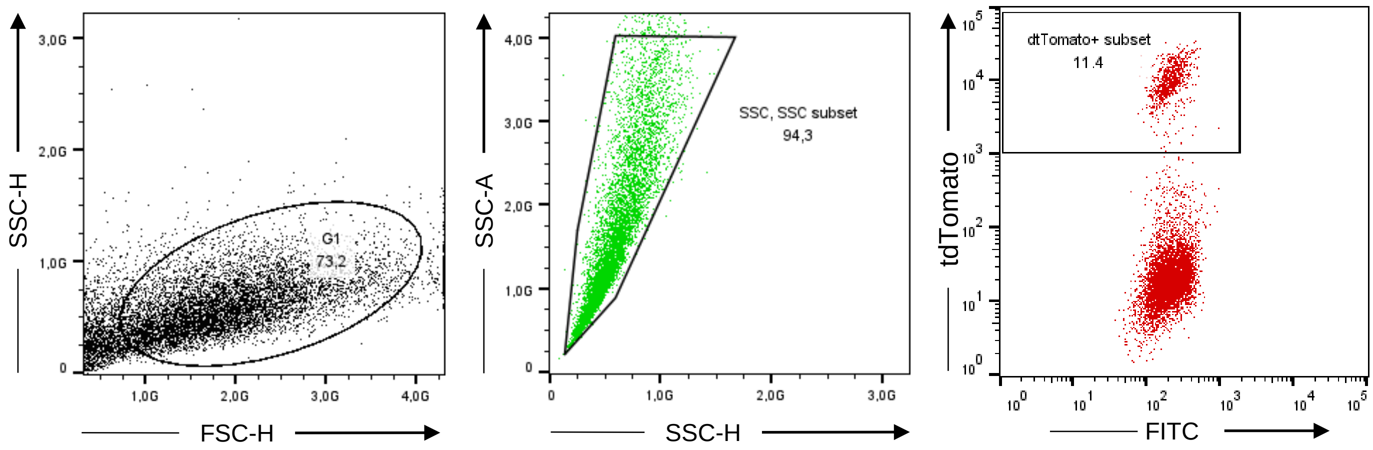


**B** Progatzky *et al.*

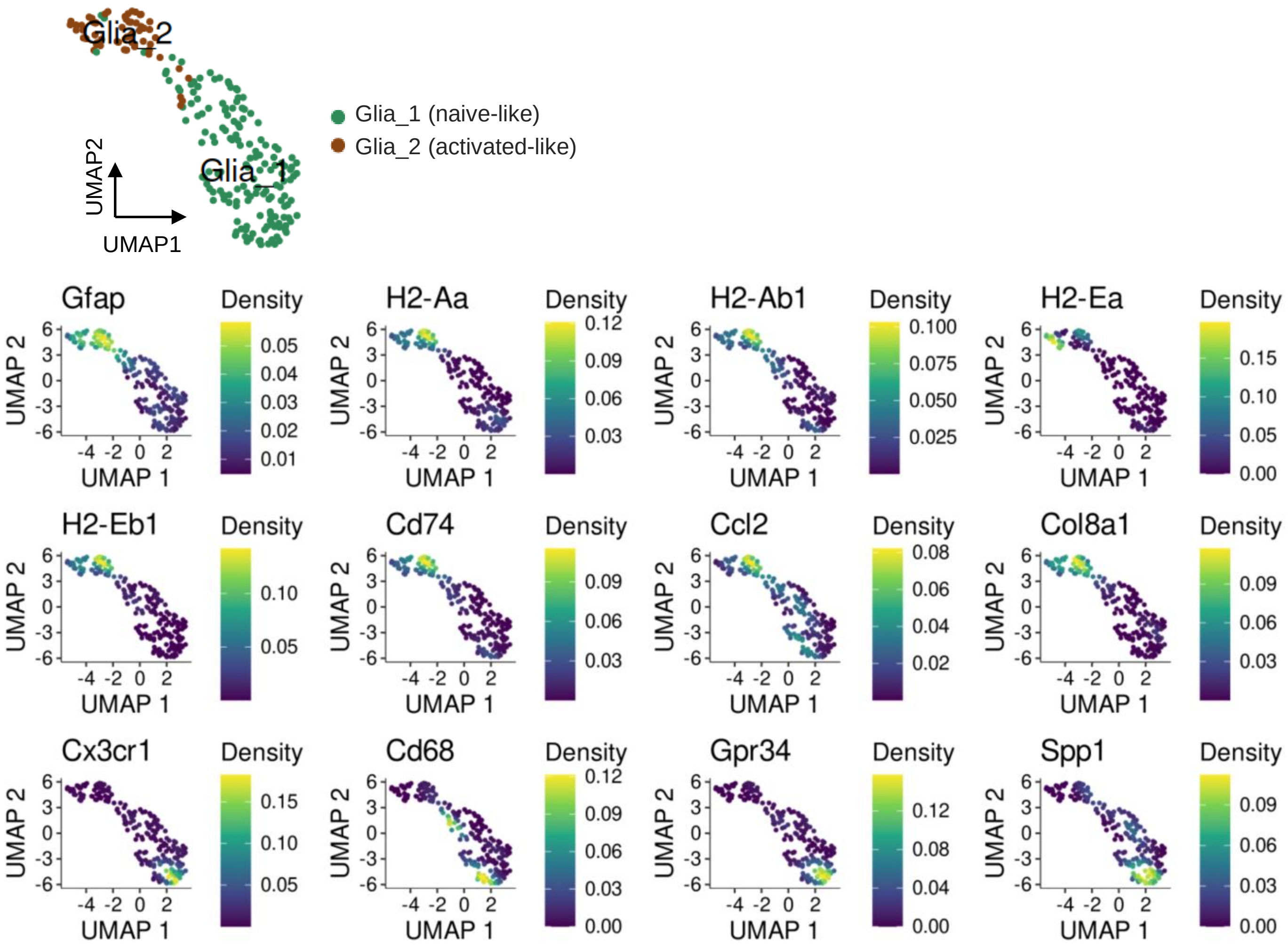

### Supplemental Figure 6

Supplemental Figure 6

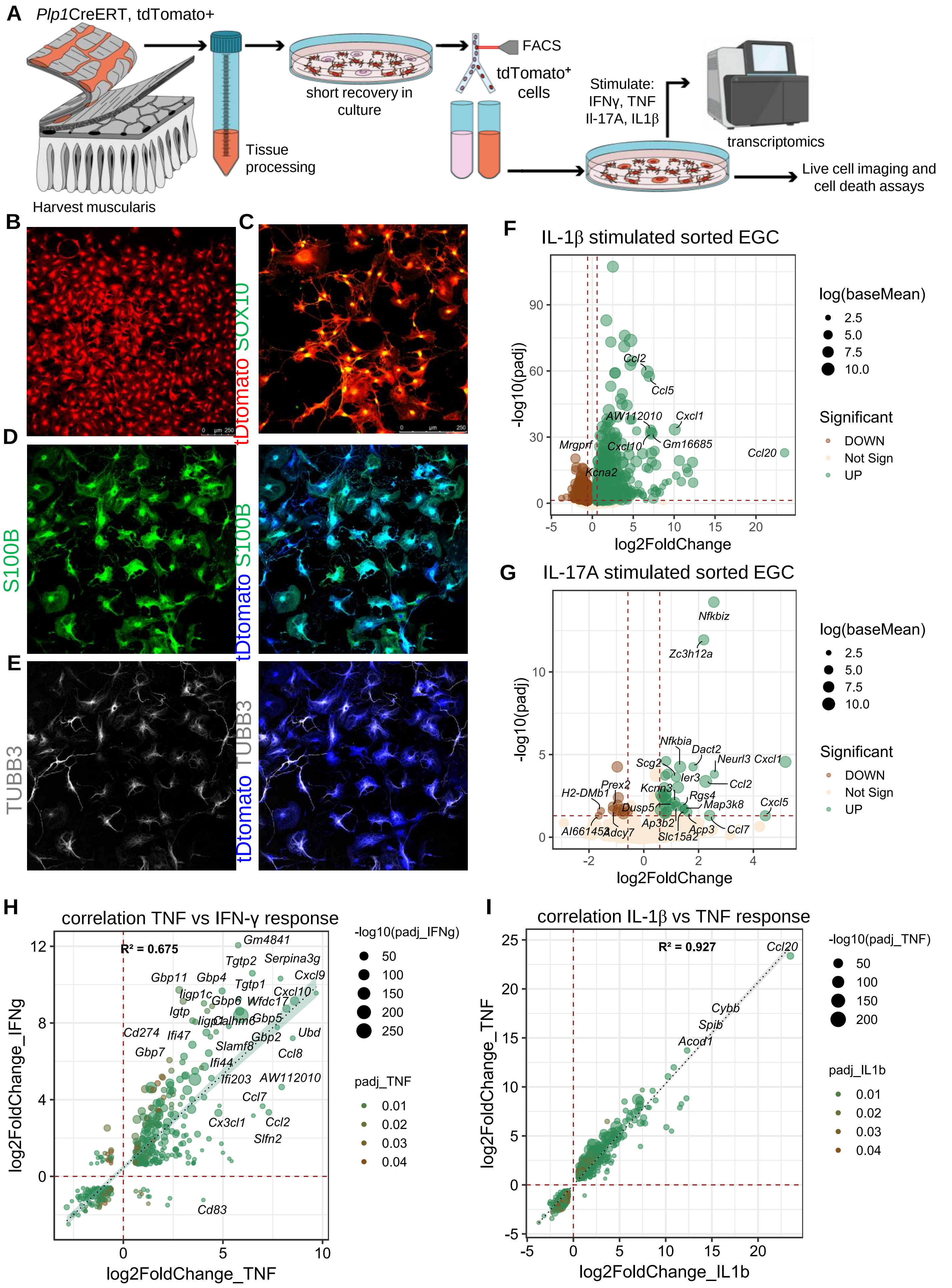

#### Supplemental Figure 6

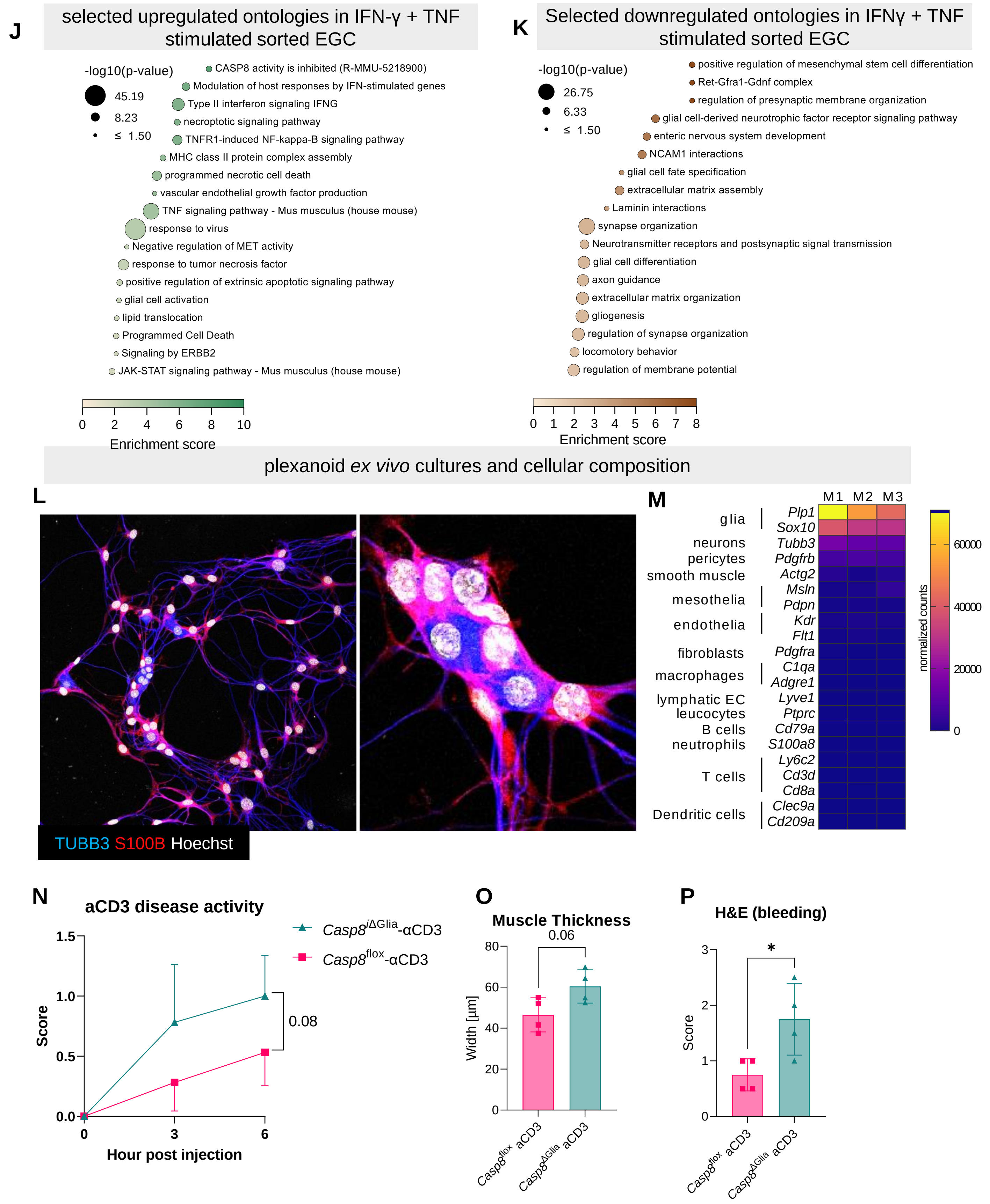
