## Supplemental Figure 5 for "Coordinated IFN-γ/TNF Axis Drives Selective Loss of Activated Enteric Glia in Inflammatory Bowel Diseases"

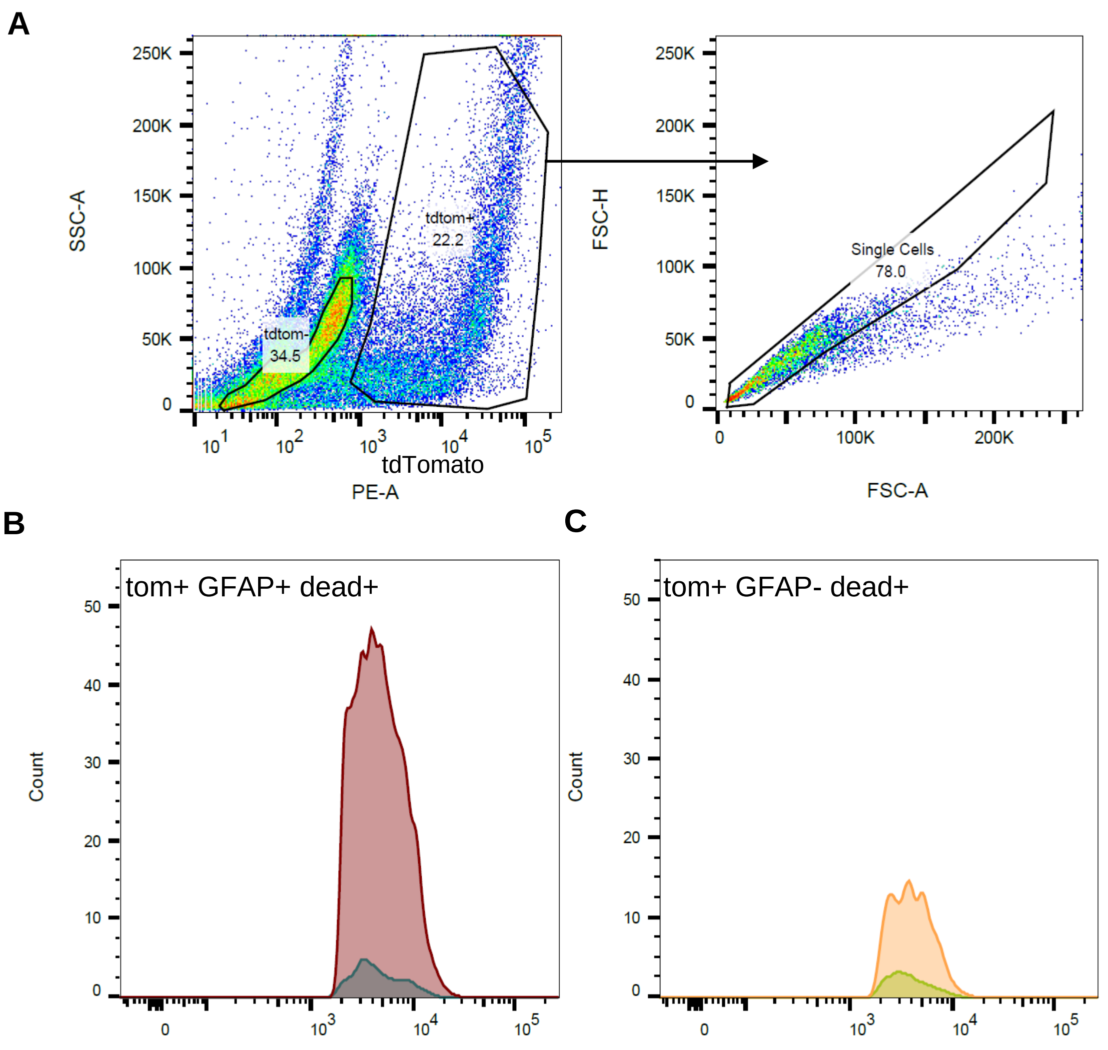


| | Sample Name | Subset Name | Count | Freq. of Total | Freq. of Parent |
|---|---|---|---|---|---|
| | 20241010_1xIsowt_1xaCD3wt_ID17_aCD3-a.fcs | Q10: Comp-APC-A+ , Comp-Pacific Blue-A+ | 1882 | 4.67 | 36.7 |
| | 20241010_1xIsowt_1xaCD3wt_ID16_Iso-a.fcs | Q10: Comp-APC-A+ , Comp-Pacific Blue-A+ | 160 | 1.55 | 14.4 |

| | Sample Name | Subset Name | Count | Freq. of Total | Freq. of Parent |
|---|---|---|---|---|---|
| | 20241010_1xIsowt_1xaCD3wt_ID17_aCD3-a.fcs | Q11: Comp-APC-A+ , Comp-Pacific Blue-A- | 488 | 1.21 | 9.52 |
| | 20241010_1xIsowt_1xaCD3wt_ID16_Iso-a.fcs | Q11: Comp-APC-A+ , Comp-Pacific Blue-A- | 100 | 0.97 | 9.00 |
